# Novel endornaviruses infecting *Phytophthora cactorum* that attenuate vegetative growth, promote sporangia formation, and confer hypervirulence to the host oomycete

**DOI:** 10.1101/2024.11.22.624937

**Authors:** Kohei Sakuta, Aori Ito, Yukiko Sassa-O’Brien, Tomohiro Yoshida, Toshiyuki Fukuhara, Seiji Uematsu, Ken Komatsu, Hiromitsu Moriyama

**Author notes:** **Correspondence:** Hiromitsu Moriyama.

## Abstract

Two novel endornaviruses were found in *Phytophthora cactorum* isolated from severe black lesions on *Boehmeria nivea* (*B. nivea*) *var*. *nipononivea* plants in a Japanese forest. These two endornaviruses were named Phytophthora cactorum alphaendornavirus 4 (PcAEV4) and Phytophthora cactorum alphaendornavirus 5 (PcAEV5), and have site-specific nick structures in their positive RNA strands, which are hallmarks of alphaendornaviruses. Ribavirin and cycloheximide treatment of the protoplasts effectively cured the host oomycete (Kara1) of the viruses. The resultant virus-free strain (Kara1-C) displayed luxuriant mycelial growth with less zoosporangia formation as compared to the Kara1 strain. Remarkably, the Kara1-C strain exhibited reduced ability to form black lesions on *B. nivea* leaves, suggesting that presence of PcAEV4 and PcAEV5 in the Kara1 strain led to enhanced virulence in host plants. Under osmotic pressure and cell wall synthesis inhibition, the Kara1 strain exhibited less growth inhibition compared with the Kara1-C strain. In contrast, the Kara1 strain showed more growth inhibition to membrane-permeable surfactant compared with the Kara1-C strain, indicating that these endornaviruses can alter the susceptibility of the host oomycete to abiotic stresses. Co-localization and cell fractionation analyses showed that PcAEV4 and PcAEV5 localized to intracellular membranes, particularly the ER membrane fraction. Further, infection with these 2 endornaviruses was found to affect the host’s response to exogenous sterols, which enhanced vegetative growth and zoosporangia formation, as well as virulence of the host oomycete. These results provide deep insights into the effects of endornavirus infection in the *Phytophthora* spp. and also highlight protoplast-based methods in advancing *Phytophthora* virus studies.

## Introduction

Mycoviruses are broadly defined as fungal and oomycete viruses, which infect the host persistently and latently, and in most cases show no obvious symptoms. Mycoviruses in fungi were first described in 1962 as viruses associated with a symptom called "die-black" in cultivated mushrooms [1]. With the discovery of Cryphonectria hypovirus 1 that infect *Cryphonectria parasitica*, which was the first mycovirus shown to cause host hypovirulence [2, 3], mycoviruses are increasingly attracting the attention of more and more researchers around the world because of their potential use in controlling plant pathogens.

Viruses that infect oomycetes belonging to the Stramenopile kingdom were first discovered in the mid-1980s by Honkura et al. in *Sclerophthora macrospora*, a causal phytopathogen of rice downy mildew disease [4]. Phytophthora alphaendornavirus 1 (PEV1), classified in the genus *Alphaendornavirus*, was the first mycovirus isolated from an oomycete of the *Phytophthora* spp. found on Douglas fir by Hacker et al. in 2005 [5]. Since then, viruses infecting oomycetes have been mainly found in *Phytophthora* spp., plant pathogens that cause devastating damages on agriculture and forests, followed by the discovery of many viruses in marine ecosystems [6, 7, 8, 9, 10]. Recently, several novel viruses that do not belong to existing virus groups have been identified by high-throughput sequencing (HTS) from *Bremia lactucae*, a downy mildew of lettuce [11]. These viruses a/re in completely novel evolutionary positions not previously found in other hosts, indicating the great diversity of viruses that infect oomycetes and the importance of further identification of them.

Several oomycete-infecting viruses are known to modify host growth. Four viruses, Phytophthora infestans RNA virus 1-4 (PiRV1-4), have been found in potato blight fungi (*Phytophthora infestans*) isolated in North America, of which PiRV2 infection stimulated sporangia formation and reduced pathogenicity [12]. Phytophthora endornavirus 2 (PEV2) and Phytophthora endornavirus 3 (PEV3), which co-infect the Japanese asparagus pathogen of the *Phytophthora* spp., inhibit host mycelial growth while increasing zoosporangia formation, alter sensitivity to pesticides, and further slow yeast cell growth when their full-length genome was introduced [9, 13]. However, as compared to fungi, the molecular biology of oomycetes is less well understood, and viral curing and viral reinfection systems have yet to be established. Due to these difficulties, the mechanisms involved in oomycete host growth and pathogenicity altered by the viruses are largely unknown.

Endornaviruses have been reported in plants, fungi, and oomycetes [5], and are classified by the International Committee on Taxonomy of Viruses (ICTV) in the family *Endornaviridae* [14]. Endornaviruses have linear plus positive-sense RNA ((+)RNA) genomes ranging from 9.8 to 17.6 kilobases (kb) in length, which contain one open reading frame (ORF) encoding a single polyprotein consisting of 3,217-5,825 amino acid (aa) residues with several conserved domains: helicase (Hel), UDP-glycosyltransferase (UGT), and RNA-dependent RNA polymerase (RdRp). A site-specific plus-strand break (nick) has also been identified as a unique genomic structure in some endornaviruses [15, 16, 17, 13]. Endornaviruses do not have an outer coat protein, but instead their replication components, dsRNAs and RdRps, are present in the endoplasmic reticulum (ER) membrane of infected cells, where their replication activity is detected [18, 19, 20]. Viruses in the order *Martellivirales*, to which the family *Endornaviridae* belongs, have been reported to remodel and replicate ER-derived membrane components, such as tobacco mosaic virus and brome mosaic virus [21, 22].

In general, (+)RNA viruses replicate within intracellular membranes, interacting with various lipids in ER, mitochondria, peroxisomes, and endosomal membranes, to encapsulate the replication intermediate, dsRNA, in the viral replicase complex (VRC) [23]. The VRC contributes to efficient genome replication by closely enveloping potential factors involved in replication and protecting them from immune defenses such as RNA silencing and interferon signaling that target dsRNA molecules [24]. Recently, sterols have been found to play an important role in VRC formation [25]. Sterols have been implicated in the meticulous packing of phospholipids and membrane stability of VRCs, and may influence the interaction of viral and host factors for replication [26]. Further, sterols in cells are also important for RNA virus replication [27, 28, 29, 30]. Peronosporales, including the genus *Phytophthora*, are sterol auxotrophs and are dependent on sterols for mycelial growth, zoosporangia formation, and zoospore production [31]. However, they do not possess genes involved in sterol biosynthesis, so they require exogenous uptake of sterols from the environment [32, 33, 31]. The involvement of intracellular membranes and sterols in the replication of (+)RNA mycoviruses, including endornaviruses, has not been investigated yet. Due to the high stability of dsRNA genomes during the extraction process, studies on mycovirus replication have been carried out exclusively on dsRNA viruses. Thus, it would be interesting to study whether (+)RNA viruses, including endornaviruses, replicate in a membrane-dependent manner and require sterols in *Phytophthora* spp., which lack a sterol biosynthetic pathway.

*Phytophthora cactorum* (*P. cactorum*) is a polyphyllous plant pathogenic oomycete with a wide host range of at least 154 genera and more than 200 species, including herbaceous and woody plants [34]. Strawberries are the most representative crop severely affected by *P. cactorum*, with seedling reductions reaching nearly 40% in Norway in 1997 and fruit losses reaching 80% in Fujian, China in 2005 [35, 36]. This pathogen has also recently been isolated from rhizosphere soils of alder (*Alnus* spp.) forests in Austria and from alder bark ulcers, and has been reported as a pathogenic oomycete in forest ecosystems [37]. In this study, two different endornaviruses were found to co-infect *P. cactorum* isolated from the perennial herbaceous plant, calamus (*B. nivea* [*B. nivea*] var. *nipononivea*), which is native to forest ecosystems in Japan, and the impact of viral infection on the host was investigated. Our findings revealed a correlation between endornavirus infection and the host fungal growth rate, zoosporangia formation, stress susceptibility, and pathogenicity. Additionally, cell fractionation and immunofluorescence staining (IFA) for dsRNA localized the endornaviruses within protoplasts, specifically associated with ER membranes. These results not only clarify the host-virus interrelationships between *Phytophthora* sp. and endornaviruses, but are expected to broaden our understanding of the dynamics of endornavirus infection in fungi, plants, and oomycete cells.

## Results

### 1. Detection and phylogenetic analysis of novel endornaviruses infecting *Phytophthora cactorum* isolated from lesions on *B. nivea* var. *nipononivea*

A dsRNA component of about 13 kb was detected in *P. cactorum* (strain Kara1) isolated from *B. nivea* var. *nipononivea* plants, which grow wild in forests in Japan and showed wilt symptoms (Fig. 1A). Illumina sequencing of total RNA extracted from the Kara1 strain generated 8,571,034 reads and 371 assembled contigs. Of these, two contigs over 12,000 base pairs (bp) were found, which were possibly related to endornavirus genomes. A BLAST search against NCBI database of the two contigs revealed that one contig showed high sequence homology (query coverage, 59%; *E*-value, 0.0; percent identity, 64.9%) with a partial sequence of Phytophthora cactorum alphaendornavirus 2 (PcAEV2), and the other contig showed high homology (query coverage, 88%; E-value, 0.0; percent identity, 74.8%) with a partial sequence of Phytophthora cactorum alphaendornavirus 3 (PcAEV3), suggesting co-infection of the Kara1 strain with two different endornaviruses. Consequently, these two contigs were tentatively designated as Phytophthora cactorum alphaendornavirus 4 (PcAEV4) and Phytophthora cactorum alphaendornavirus 5 (PcAEV5), respectively, because of their homology to PcAEV2 and PcAEV3 and were thus named sequentially. Based on the sequences of both contigs, specific primer sets were designed to distinguish these two endornaviruses. Two bands were detected by duplex RT-PCR using these two primer sets, confirming that the Kara1 strain was co-infected with PcAEV4 and PcAEV5 (Fig. 1B). To determine the 5’- and 3’-terminal sequences of the genomes, 5’ and 3’ RACE were performed, which revealed the complete genomes to be 12,839 nucleotides [nt] (PcAEV4) and 12,774 nt (PcAEV5) in length. Both sequences were deposited in the GenBank database under accession numbers LC847181(PcAEV4) and LC847182 (PcAEV5). The PcAEV4 genome is consisted of a 168 nt 5′ untranslated region (UTR), a single ORF consisting of 4198 aa, and a 74 nt 3′ UTR; the PcAEV5 genome is consisted of a 134 nt 5′ UTR, a single ORF consisting of 4196 aa, and a 49 nt 3′ UTR.

**Figure 1.**
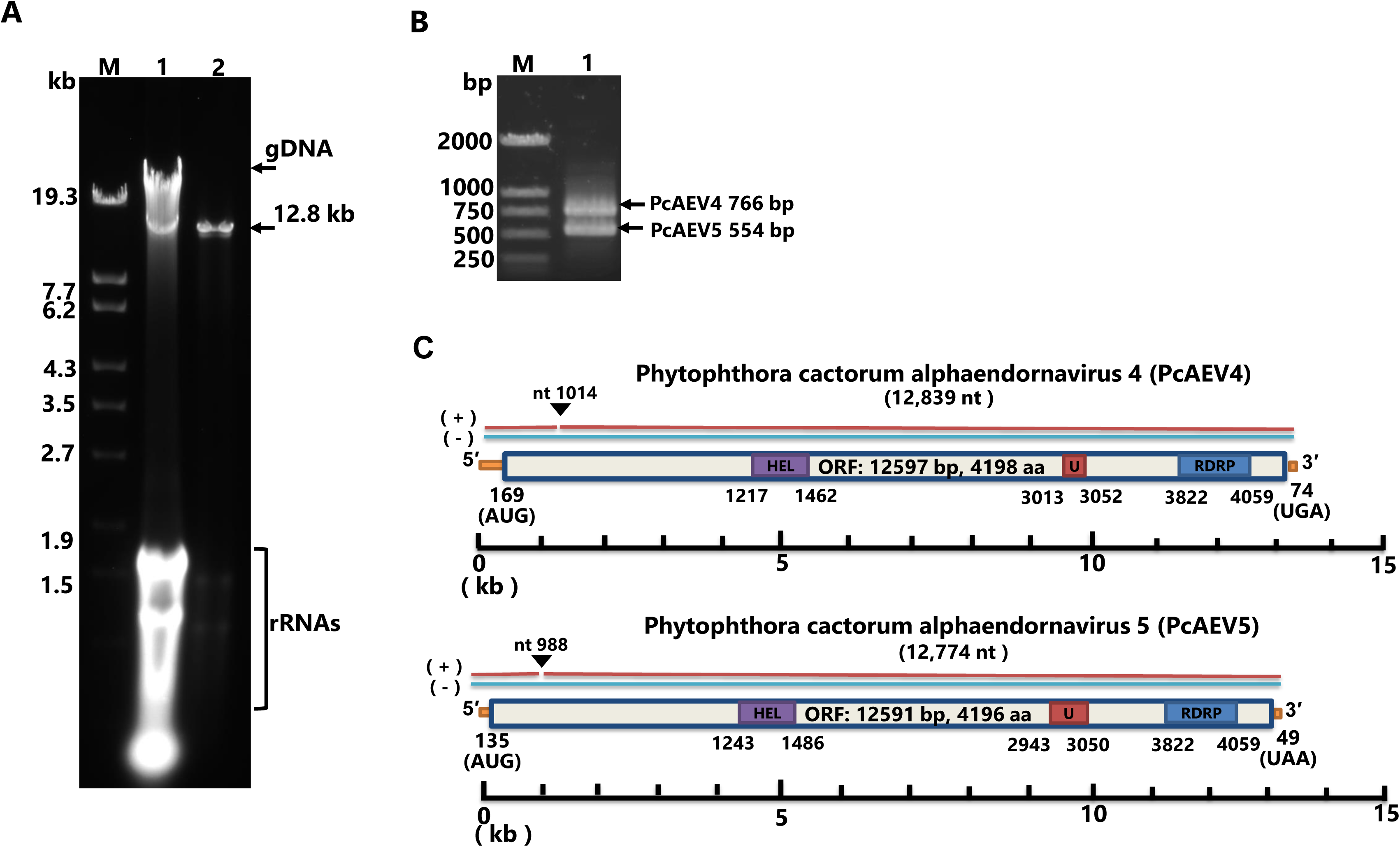
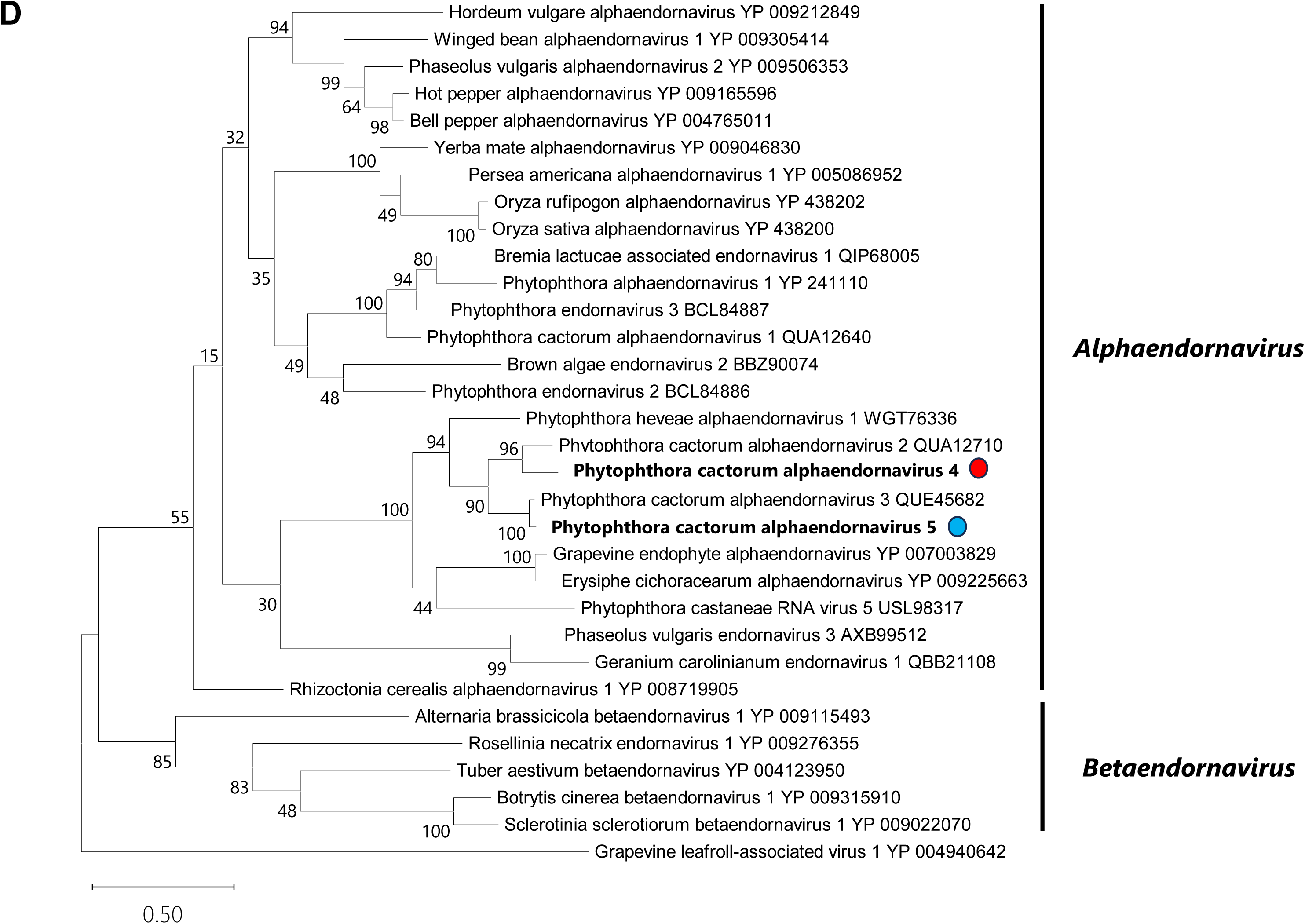
Detection of PcAEV4 and PcAEV5 in the Kara1 strain of *P. cactorum*. **(A)** Agarose gel electrophoresis of dsRNA purified from the Kara1 strain. Lane designations: M: DNA marker (250 ng of λ DNA digested with EcoT14I); 1, total nucleic acid; 2, purified dsRNA. Electrophoresis was performed on 0.8% agarose gel at 40V for 6 h, followed by staining with ethidium bromide (0.5 μg/ml). The arrows indicate the position of the 12.8 kb dsRNA and genomic DNA (gDNA). **(B)** Simultaneous detection by duplex RT-PCR using PcAEV4- and PcAEV5-specific primers. Lane designations: M, DNA size marker; 1, dsRNA extracted from Kara1. **(C)** Genomic maps of PcAEV4 and PcAEV5 showing predicted amino acid numbers. Boxes represent large ORFs, and lines represent UTRs. HEL, viral helicase 1; U, UDP-glycosyltransferase; RdRp, viral RNA-dependent RNA polymerase. **(D)** Maximum likelihood tree (RAxML) showing the phylogenetic relationships of the putative RdRp of endornaviruses. Nodes are displayed with bootstrap support values ≥50%. Branch lengths are scaled to the expected number of amino acid substitutions per site. GenBank accession numbers of the analyzed genes are provided in Supplemental Table 1. Grapevine leafroll-associated virus 1, an Ampelovirus of the family *Closteroviridae*, was used as an outgroup.

NCBI BLAST search using the full-length predicted amino acid sequences of PcAEV4 and PcAEV5 as queries revealed the presence of three domains: a putative RNA helicase 1 (Hel-1) (PcAEV4: 1217-1462 aa, *E*-value = 3e-09; PcAEV5: 1243-1369 aa, *E*-value = 4e-04), UGT (PcAEV4: 3013-3052 aa, *E*-value = 0.016; PcAEV5: 2943-3050 aa, *E*-value = 0.004), and RdRp (PcAEV4: 3822-4059 aa, *E*-value = 2e-88; PcAEV5: 3822-4059 aa, *E*-value = 7e-88) (Fig. 1C). The amino acid sequence of PcAEV4 was the most similar (query coverage, 99.0%; *E*-value, 0.0; percent identity, 58.6%) to the predicted polyprotein sequence of PcAEV2 (QUA12642), and PcAEV5 was found to have 84.4% identity (query coverage, 99.0%; *E*-value, 0.0) to the predicted protein sequence of the closely related PcAEV3. Next, a phylogenetic tree based on the maximum likelihood method was constructed using the estimated RdRp regions of PcAEV4, PcAEV5, and 30 other endornaviruses, with Grapevine leafroll-associated virus 1 as an outgroup. Phylogenetic analysis showed that PcAEV4 and PcAEV5 belong to the genus *Alphaendornavirus*, and form a monophyletic cluster with Phytophthora heveae alphaendornavirus 1, PcAEV2, and PcAEV3, which is distinct from PcAEV1 (Fig. 1D). Pairwise comparisons of the nucleotide and amino acid sequences of PcAEV1, PcAEV2, PcAEV3 [8], PcAEV4, and PcAEV5 showed that the genomes of these five viruses exhibit 44.3-73.2% nucleotide and 19.2-83.7% amino acid sequence identity, with 32.5-91.9% amino acid similarity (Fig. S1). PcAEV1 showed low sequence homology to the other four endornaviruses from the same host fungus, which supports its placement in a different phylogenetic cluster. PcAEV3 and PcAEV5 are the most similar to each other among these five viruses, which showed the highest nucleotide sequence identity (73.2%), with even higher amino acid sequence similarity (91.9 %). These results suggested genetic diversity among endornavirus species infecting *P. cactorum* with different habitats and host plants. According to the species demarcation criteria for the genus *Alphaendornavirus* established by the ICTV (same species when more than 75.0% overall sequence identity), PcAEV4 and PcAEV5 should be considered as a new species in the family *Alphaendornaviridae*.

### 2. Detection of a nick in PcAEV4 and PcAEV5 genomes

The novel endornaviruses PcAEV4 and PcAEV5 identified in the Kara1 strain were examined for the presence of a nick, a genomic structure characteristic of alphaendornaviruses. Two double-stranded DNA probes were designed to anneal to the 5’ and 3’ terminal sequences of the genomes of PcAEV4 and PcAEV5 (Fig. 2A, 2B). Northern blot analysis showed that two bands of approximately 12 kb representing the entire genomes of PcAEV4 and PcAEV5 were detected by all four probes, while a band of approximately 1 kb representing the nick was detected exclusively by the PcAEV4-5’ and PcAEV5-5’ probes (Fig. 2C, 2D, 5’ probe). Using the PcAEV4-3’ and PcAEV5-3’ probes, two bands around 12 kb were detected (Fig. 2C, 2D, 3’ probe). This may indicate the 1 kb difference between the positive and negative strands created by the nick, suggesting that the upper band was the negative strand and the lower band was the positive strand. These results indicated that PcAEV4 and PcAEV5, which co-infect Kara1, have a nick in the 5’ terminal region.

**Figure 2.**
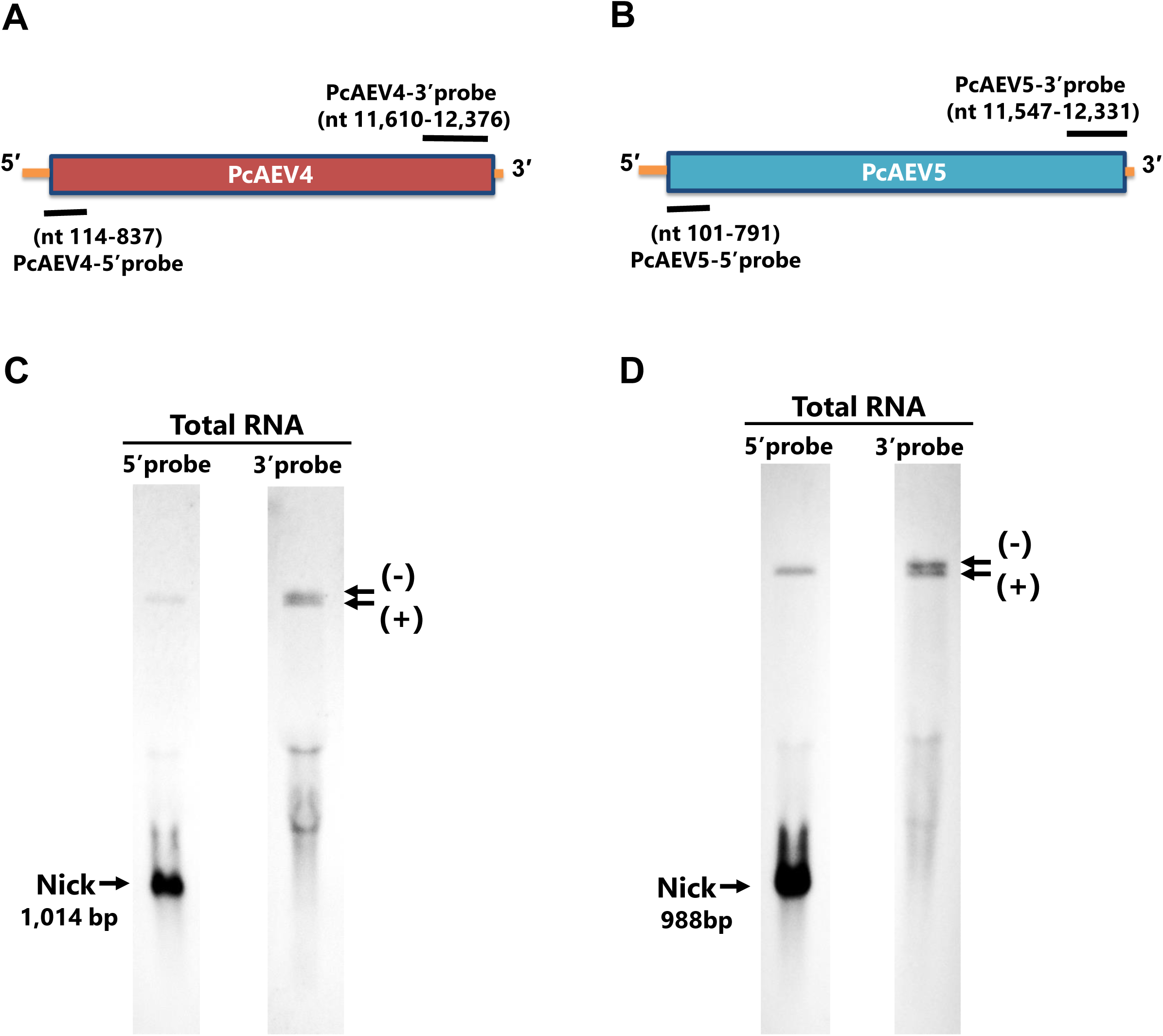
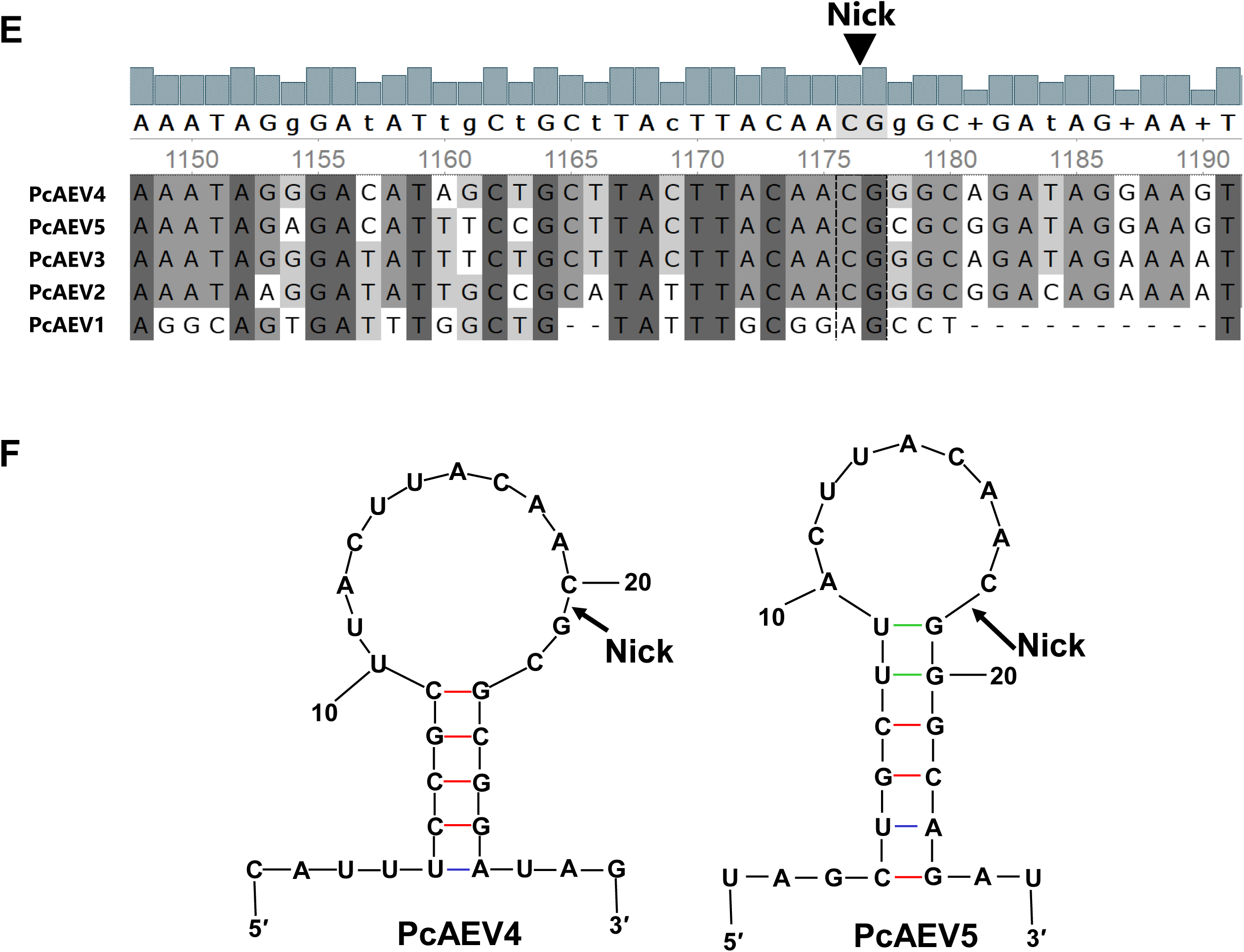
Identification and sequencing of nicks in PcAEV4 and PcAEV5 co-infecting the Kara1 strain via Northern blot analysis. Schematic genomic maps of PcAEV4 **(A)** and PcAEV5 **(B)**. Northern blot hybridization to detect nicks in PcAEV4 **(C)** and PcAEV5 **(D)**. Approximately 20 µg of total RNA extracted from the Kara1 strain was loaded, and DIG-labeled DNA probes were used for detection. The probes were prepared at the following positions: PcAEV4-5’(nt 114-837), PcAEV4-3’(nt 11,610-12,037), PcAEV5-5’(nt 101-791), and PcAEV5-3’ (nt 11,547-12,331). **(E)** Multiple sequence alignment of the nucleotide sequences surrounding the nicks identified in the endornaviruses of *P. cactorum*. The alignment was performed using CLUSTALW, with perfectly matched nucleotides shaded in black and partially matched nucleotides shaded in gray. **(F)** Secondary structure prediction of the nick sequences of PcAEV4 and PcAEV5. Secondary structure prediction was carried out using the mfold server [63]. Black arrows indicate the cleavage sites of the nicks.

To determine detailed nick locations, 3’ RACE was performed. Sequence analysis showed that PcAEV4 and PcAEV5 have a 3’ terminal poly(A) tail at nt 988 and 1,013, respectively, suggesting that the sequence up to this cleavage point is nicked. Comparison of the sequences around the nick in the five *P. cactorum* endornaviruses by multiple alignments using CLUSTALW showed eight identical nucleotides (TTACAAC/G: / denotes cleavage site) in four endornaviruses except PcAEV1 (Fig. 2E). RNA secondary structure prediction of sequences surrounding the nick position by Mx fold [38] showed that both PcAEV4 and PcAEV5 form stem-loops, including a 15-base loop structure, which includes the nick position (Fig. 2F). The presence of a nick in PcAEV2 and PcAEV3 has not yet been investigated, but the high conservation of these regions suggests that these two viruses may also have a similar nick.

### 3. Isolation of PcAEV4- and PeAEV5-cured strains

To determine the effect of PcAEV4 and PcAEV5 co-infection on the host Kara1 strain, we attempted to isolate virus-cured strains by adding the antiviral agent, ribavirin, in combination with the protein synthesis inhibitor, cycloheximide, to protoplasts prepared from the virus-infected strain. After isolating 16 strains from the regenerated protoplasts, dsRNA was purified from each strain and viral infection was examined by dsRNA electrophoresis and RT-PCR using primers that specifically detect PcAEV4 and PcAEV5. PcAEV4 and PcAEV5 RNA was also not detected in any of the 16 strains neither by dsRNA electrophoresis (Fig. 3A) nor by RT-PCR (Fig. 3B). These results showed that PcAEV4 and PcAEV5 were cured in these 16 regenerated strains, indicating that the combination treatment of ribavirin and cycloheximide in protoplasts is effective in curing endornaviruses in *Phytophthora* spp. We chose one of these 16 virus-cured strains for further experimentation, hereafter called Kara1-C.

**Figure 3.**
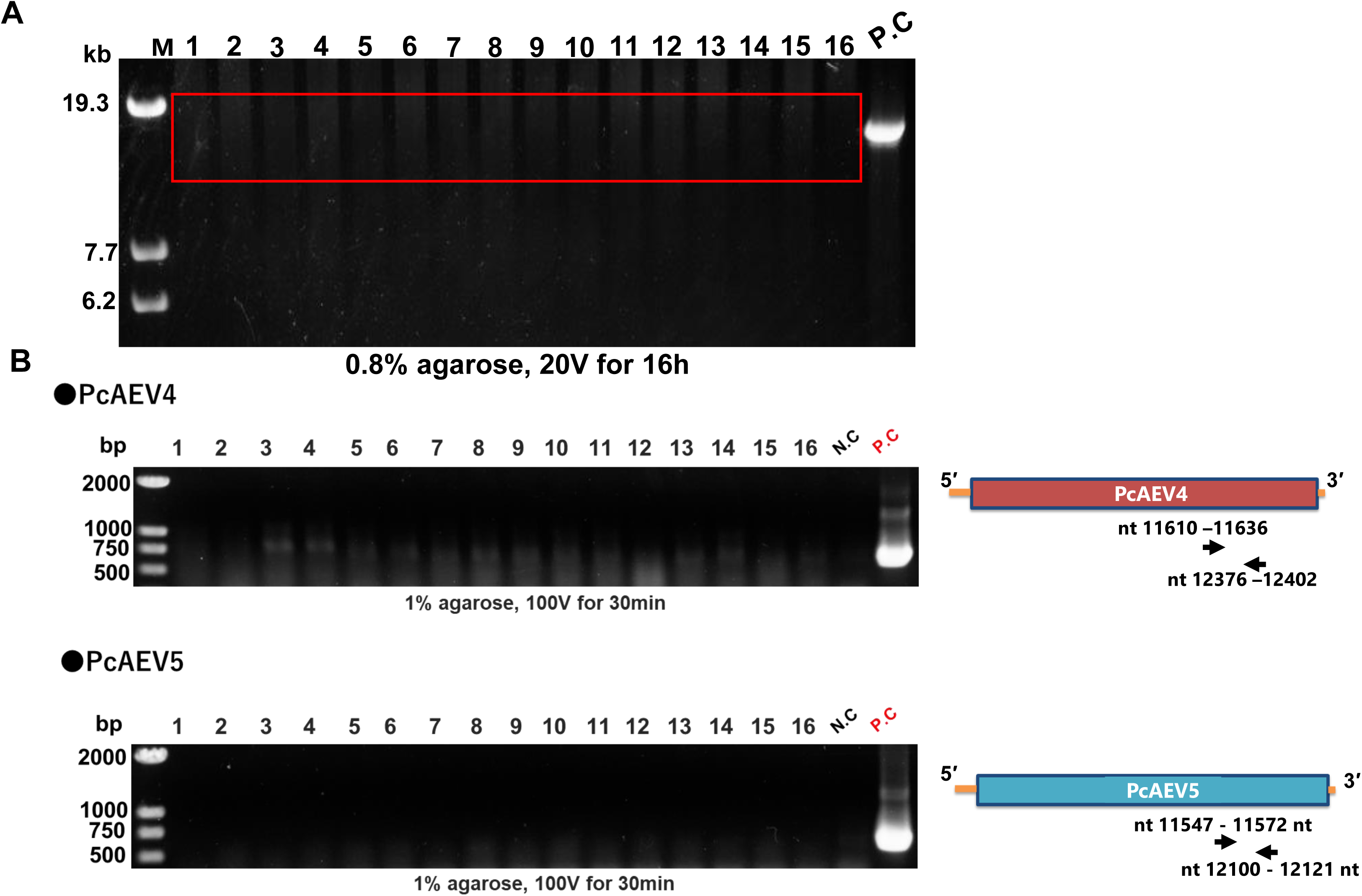
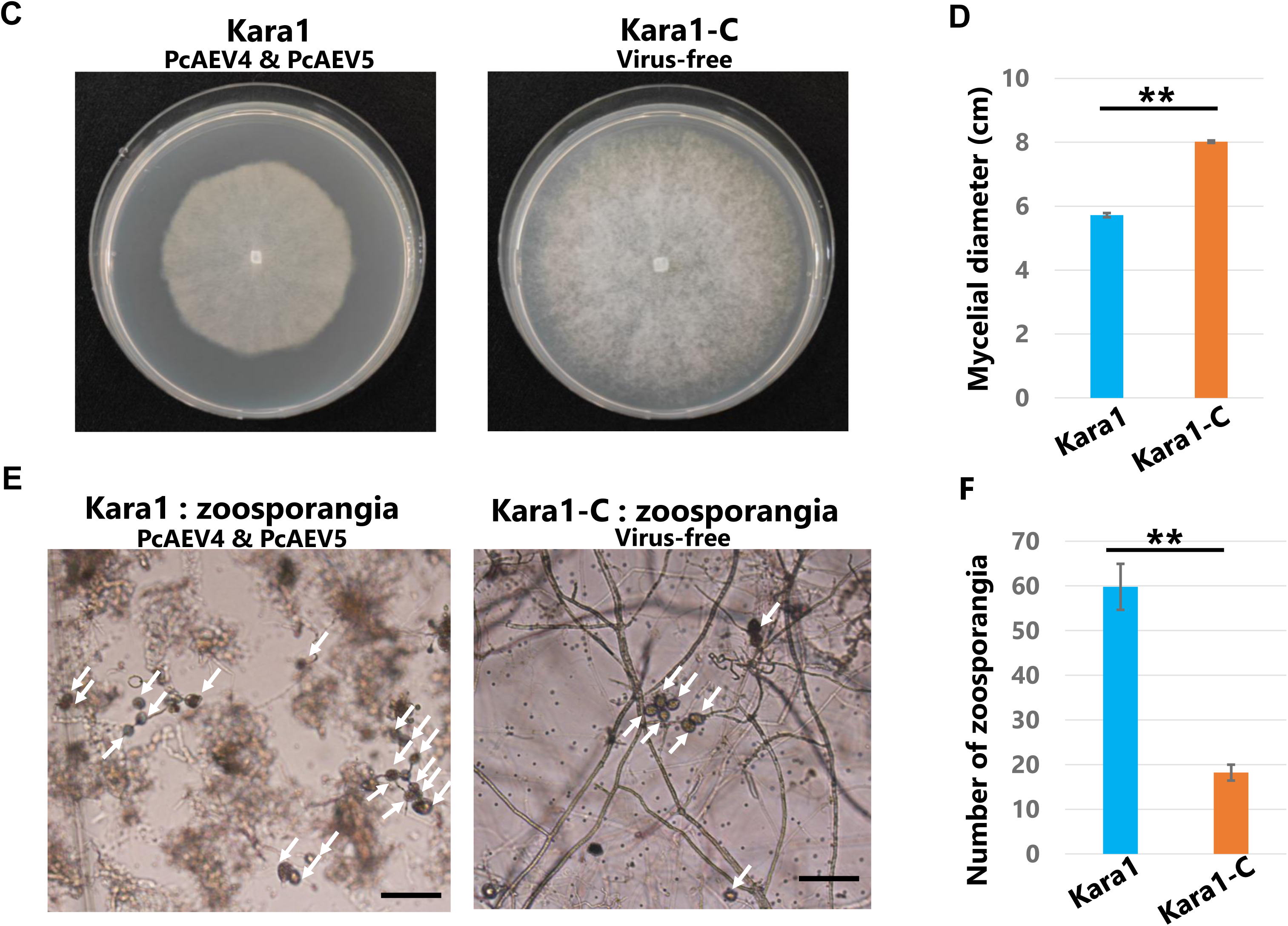
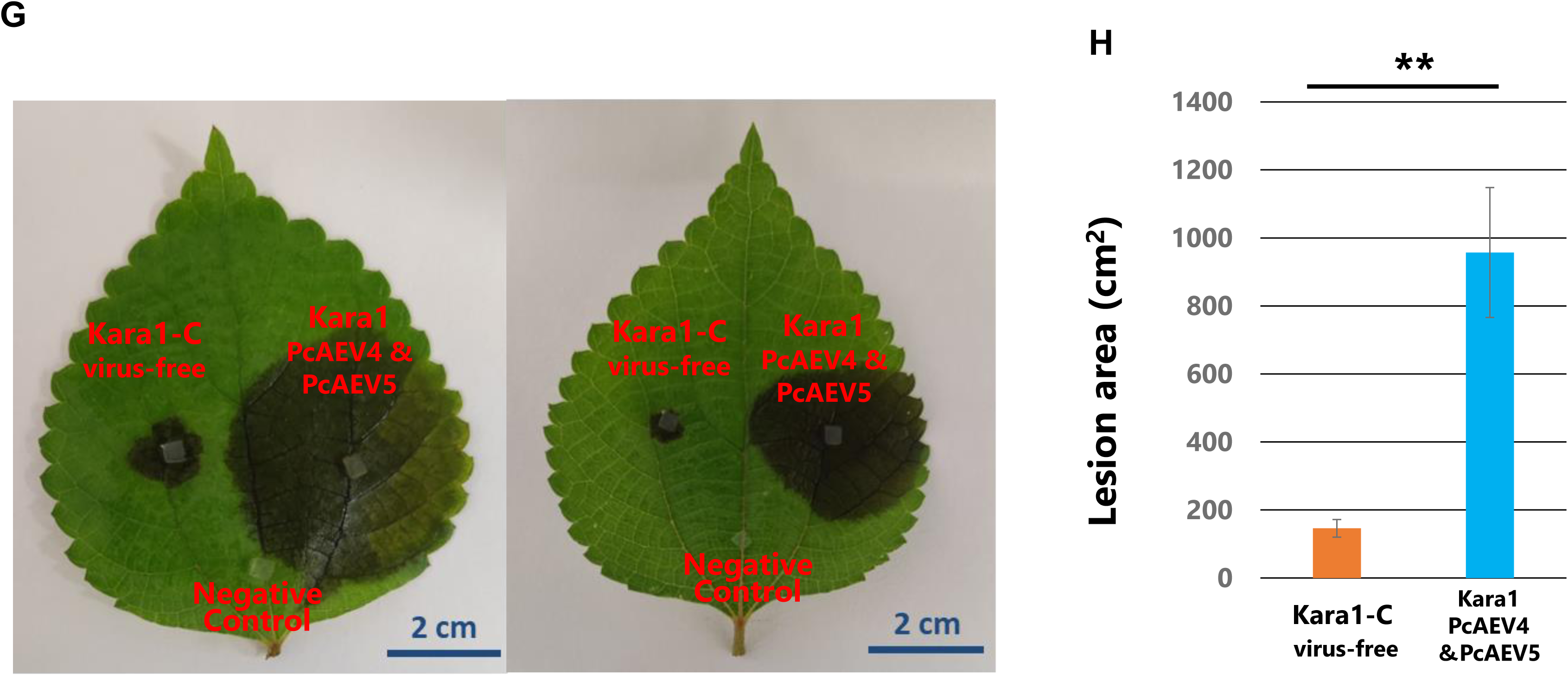
Screening of virus-cured strains and comparison of phenotypes between the Kara1 and Kara1-C strains. **(A)** Electrophoresis of purified dsRNA extracted from protoplast-regenerated strains treated with ribavirin and cycloheximide. Electrophoresis was performed on 0.8% agarose gel at 20V for 16 h. P.C: Positive control, purified dsRNA extracted from the Kara1 strain. **(B)** Confirmation of PcAEV4 and PcAEV5 infection by one-step RT-PCR. RT-PCR was performed using purified dsRNA as a template with the following primer sets: PcAEV4-3’ (nt 11,610-12,037) and PcAEV5-3’(nt 11,547-12,331) probes. N.C: Negative control, distilled water. P.C: Positive control, RT-PCR performed using dsRNA extracted from the Kara1 strain as template. **(C)** Representative images showing the morphologies and **(D)** histogram that compares mycelial diameter of the colonies of the Kara1 and Kara1-C strains cultured on V8A medium for 7 days. **(E)** Representative images showing the morphologies and **(F)** histogram that compares number of zoosporangia formation on cellophane membranes placed on V8A medium. Zoosporangia induction was performed by incubating hyphae cultured on cellophane membranes in distilled water under light conditions for 40 hours, and zoosporangia in randomly selected fields were counted. **(G)** Representative images showing lesions and **(H)** histogram that compares lesion area induced by the Kara1 and Kara1-C strains on the detached leaves of *B. nivea* var. *nipononivea*. Photographs were taken 2 days post-inoculation. Pathogenicity differences were quantified by measuring lesion areas on *B. nivea* var. *nipononivea* leaves at 2 dpi using ImageJ. Negative control was inoculation with distilled water on a different part of the same leaf. Error bars represent standard deviations from 5-6 biological replicates of a representative experiment. Statistical analysis was performed using Student’s t-test (**P < 0.01).

To investigate the effect of PcAEV4 and PcAEV5 co-infection on host growth, we compared mycelial growth on V8A medium between the PcAEV4- and PcAEV5-infected Kara1 strain and the virus-cured Kara1-C. Kara1-C showed significantly faster mycelial growth and more vigorous aerial mycelium formation (P<0.01) (Fig. 3C-3F). In contrast, the number of sporangia formed on the cellophane membrane was significantly higher in the Kara1 strain infected with PcAEV4 and PcAEV5 compared with Kara1-C (P<0.01) (Fig. 3D, 3F). To determine whether PcAEV4 and PcAEV5 co-infection affect the virulence of host *P. cactorum*, pathogenicity tests were conducted on leaves of *B. nivea* var. *nipononivea*. While both strains infected *B. nivea* var. *nipononivea* leaves and formed lesions, they were larger when produced by the virus-infected Kara1 strain (P<0.01), thus suggesting increased virulence (Fig. 3G, 3H). These results indicated that viral curing of the host *P. cactorum* significantly increased mycelial growth on V8A medium (P<0.01). However, the ability to form sporangia and lesion formation in plants were decreased in the virus-cured Kara1-C strain, suggesting that the endornaviruses increased virulence of the oomycete pathogen to the plants.

### 4. PcAEV4 and PcAEV5 co-infection alters the susceptibility of the host oomycete to different stresses

Comparison of mycelial growth after 7 days of incubation with different temperatures on V8A medium showed that there was no significant difference in mycelial size between the virus-infected and virus-cured strains between 4°C and 16°C (Fig. 4A, 4B). At 20°C (Kara1-C, 6.73 cm; Kara1, 5.63 cm) and 25°C (Kara1-C, 7.83 cm; Kara1, 6.53 cm) (P<0.01), the optimal growth temperature for *P. cactorum*, the Kara1-C strain showed significantly faster mycelial growth. At 30°C, a high temperature for *P. cactorum*, mycelial growth was observed for Kara1, but not for Kara1-C. Taken together, these results indicated that there were no observed differences in growth between Kara1 and Kara1-C strains under low temperature conditions. However, a difference in growth was found between the two strains under higher temperatures (20-30°C).

**Figure 4.**
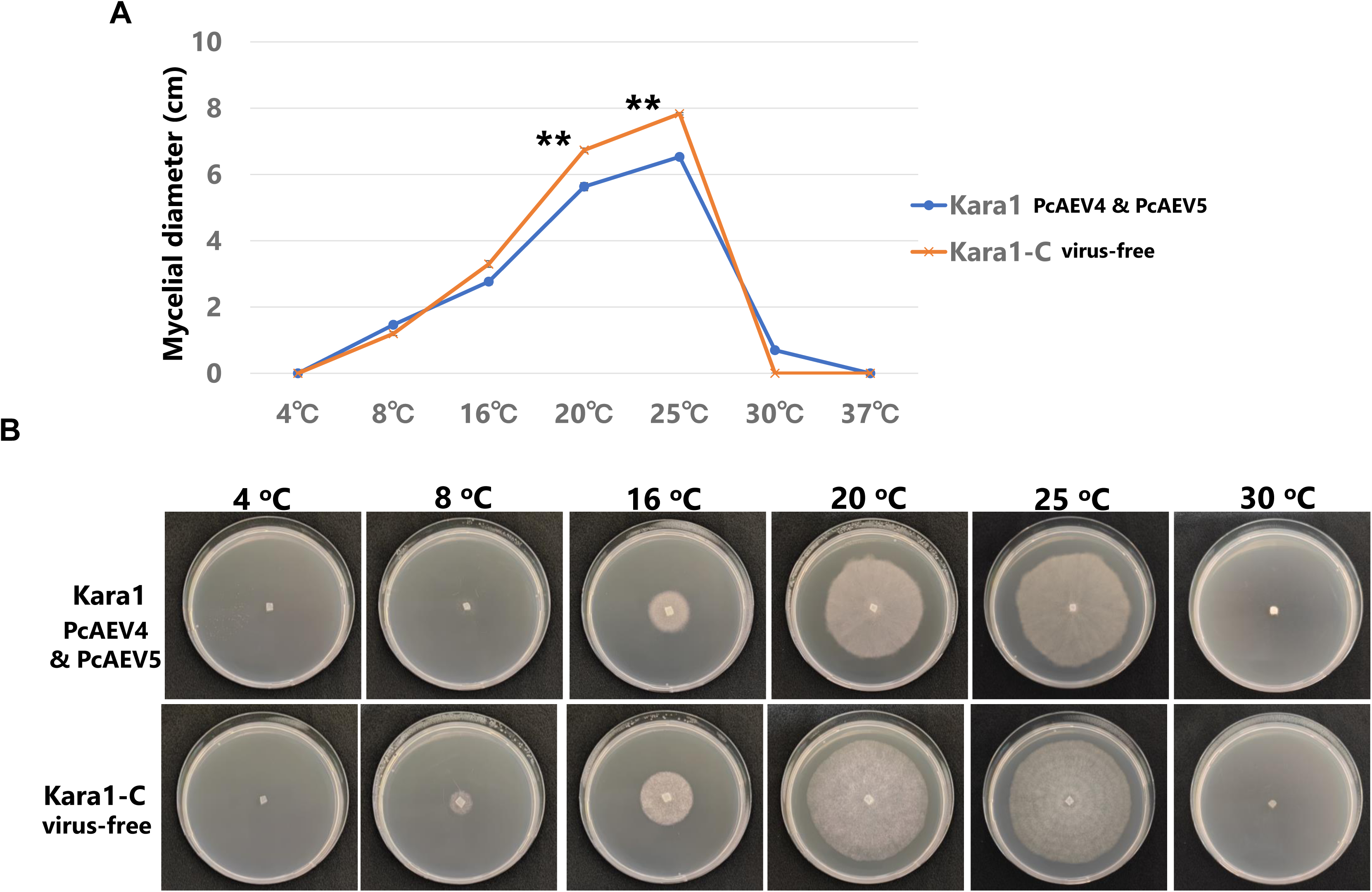
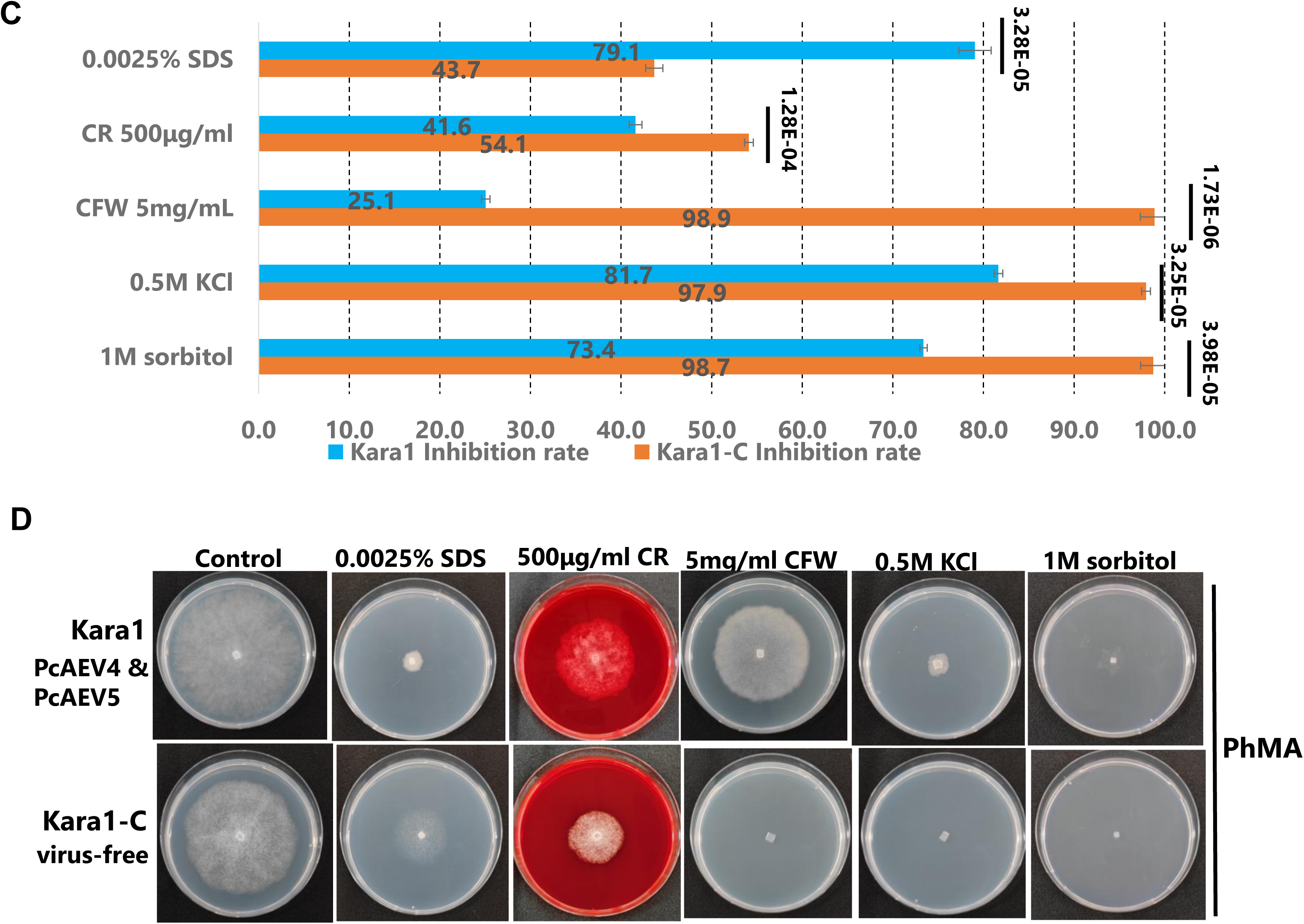
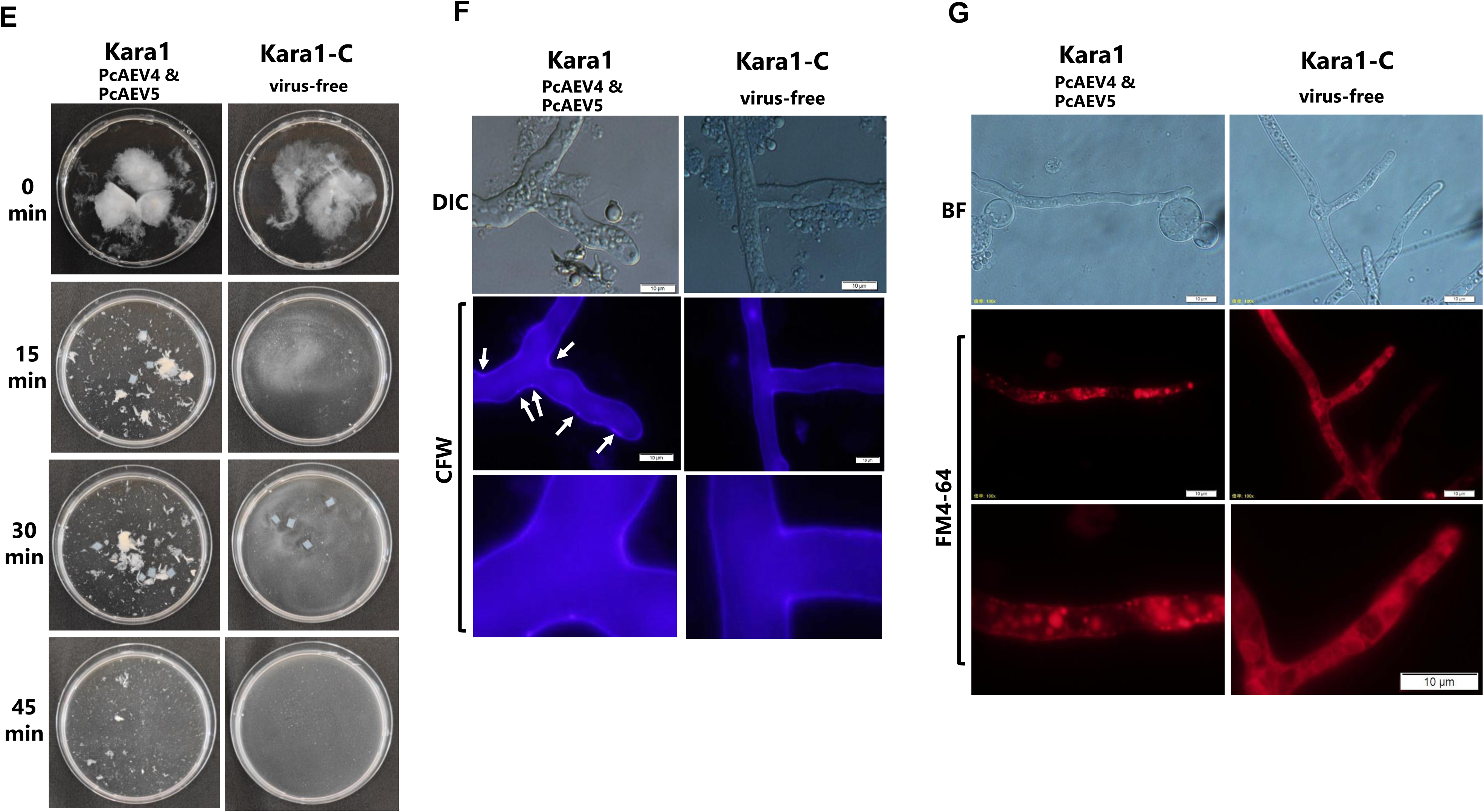
Comparison of growth of the Kara1 and Kara1-C strains under various stress conditions. **(A)** Histogram comparing the average colony diameters and **(B)** representative colony morphologies of the Kara1 and Kara1-C strains grown on V8A medium at different temperatures (4°C, 8°C, 16°C, 20°C, 25°C, 30°C, 37°C) for 7 days. **(C)** Comparison of stress sensitivity between the Kara1 and Kara1-C strains. The blue bars represent the results for the Kara1 strain, and the orange bars represent the Kara1-C strain. The values inside the bars indicate the inhibition rates (%). Error bars represent standard deviations from three biological replicates of a representative experiment. Statistical analysis was conducted using Student’s t-test, with the p-values indicated to the right of the black bars. **(D)** Representative colony morphologies of the Kara1 and Kara1-C strains grown for 10 days on PhMA medium exposed to different stress-inducing agents. **(E)** Comparison of cell wall integrity between the Kara1 and Kara1-C strains. Hyphal morphologies were observed after 15, 30, and 45 minutes of incubation in an isotonic solution containing cellulase and lysing enzyme. Equal amounts of hyphae from each strain were used for each treatment. **(F)** Calcofluor White (CFW) and **(G)** FM4-64 staining and visualization by fluorescence microscopy of hyphae from the Kara1 and Kara1-C strains. White arrows indicate strongly stained regions. Scale bar = 10 μm.

To compare the susceptibility of Kara1 and Kara1-C strains to abiotic stresses, 0.5 M KCl and 1 M sorbitol, both of which induce osmotic pressure, as well as cell wall synthesis inhibitors (Calcofluor white: CFW and congo red: CR) and membrane-permeable surfactant sodium dodecyl sulfate (SDS) were used to assess fungal growth on Phytophthora minimal agar (PhMA) medium. Growth inhibition was calculated by measuring the mycelial diameter of the Kara1 and Kara1-C strains 10 days after inoculation and comparing it to that of the control medium, in which no stress-inducing substances were added. Under hyperosmotic conditions by addition of 0.5 M KCl or 1.0 M sorbitol, inhibition of growth in the Kara1-C strain was 97.9% and 98.7%, respectively, which were significantly higher than 81.7% and 73.4% in the Kara1 strain (P<0.01) (Fig. 4C, 4D). An interesting result was that the inhibitory effect of CFW, an inhibitor of cell wall synthesis, on fungal growth was dramatically higher in the Kara1-C strain (98.9%) as compared to the Kara1 strain co-infected with PcAEV4 and PcAEV5 (25.1%) (Fig. 4C, 4D). The Kara1-C strain showed 54.1% growth inhibition by CR, whereas it was lower in the Kara1 strain (41.6%) (Fig. 4C, 4D). On the other hand, growth inhibition by SDS was much higher in the Kara1 strain (79.1%) compared with Kara1-C strain (43.7%), suggesting that the PeAEV4 and PeAEV5 co-infected strain is highly sensitive to membrane-permeable surfactants (Fig. 4C, 4D).

Since membrane stability may be altered by endornavirus infection, we next investigated the effects of co-infection with PcAEV4 and PcAEV5 on lipid polarity and cell wall biosynthesis by evaluation of protoplast reformation. We compared the time required for hyphae digestion and staining of regenerated hyphae using CFW for visualization of cellulose accumulation and FM4-64 for lipid membrane staining between the Kara1 and Kara1-C strains. Mycelia were collected in equal weight after incubation on V8A liquid medium, and an enzyme solution (cellulase: 0.75%, lysing enzyme: 0.6%, final concentration) was added to the mycelia to observe the time-dependent digestion of the cell wall. In the Kara1-C strain, almost all mycelium was digested after 15 min, whereas mycelial fragments were still visible even after 45 min in the Kara1 strain, indicating that protoplastization was delayed (Fig. 4E). After the protoplasts were incubated overnight on regeneration medium, the newly-formed cell walls were stained with CFW and observed under fluorescence microscopy. In the Kara1 strain, strong deposition of CFW could be observed in several areas, but no clear deposition could be observed in the Kara1-C strain. This result indicated that the Kara1-C strain has reduced cellulose biosynthetic capacity, since CFW binds strongly to cellulose, one of the cell wall components (Fig. 4F). These results suggested that cell wall of the Kara1-C strain is weakened, supporting the results of the CFW sensitivity test, in which the Kara1-C strain showed increased growth inhibition (P<0.01) (Fig. 4C, 4D).

We further used FM4-64 staining, which stains cell membranes and endocytic vesicles, in combination with fluorescence imaging of the protoplast-regenerated mycelia since it is well known that viral infections can induce an increase and redistribution of lipid droplets within host cells (Boulant et al., 2008; Dansako et al., 2014). Therefore, it was hypothesized that co-infection with PcAEV4 and PcAEV5 may lead to differences in lipid membrane localization and polarity between the Kara1 and Kara1-C strains. Indeed, in the Kara1 strain, there were many unusually large red granules in the mycelium, and the staining pattern was uneven (Fig. 4G, top). In contrast, the entire mycelium of Kara1-C was stained uniformly, and the outline of the vacuoles was clearly observed (Fig. 4G, lower panel). These results suggested that PcAEV4 and PcAEV5 co-infection may disrupt the lipid membrane arrangement in the host cell, which may have contributed to the higher sensitivity of the Kara1 strain to SDS (membrane-permeable surfactant) than the Kara1-C strain.

### 5. Virulence of PcAEV4- and PcAEV5-infected strain is increased by addition of sterols

Since sterols are important for both RNA virus replication and *Phytophthora* spp. growth, we used PhMA, a synthetic medium completely deficient in sterols, with or without β-sitosterol (25 µg/ml) to determine the effect of sterols on host growth/differentiation and endornavirus infection in the Kara1 and Kara1-C strains. PhMA was chosen for the following experiments because the V8A medium contains small amounts of sterols. Kara1 and Kara1-C strains were passaged twice on PhMA medium prior to measurement in order to eliminate the influence of residual sterols in the cells. After 7 days of culture at 25°C on PhMA medium, the mycelial growth of the Kara1-C strain was significantly reduced than that of the Kara1 strain (Fig. 5A). There was no difference in growth between the Kara1 and Kara1-C strains grown on PhMA medium containing β-sitosterol (PhMA+S) (Fig. 5B).

**Figure 5.**
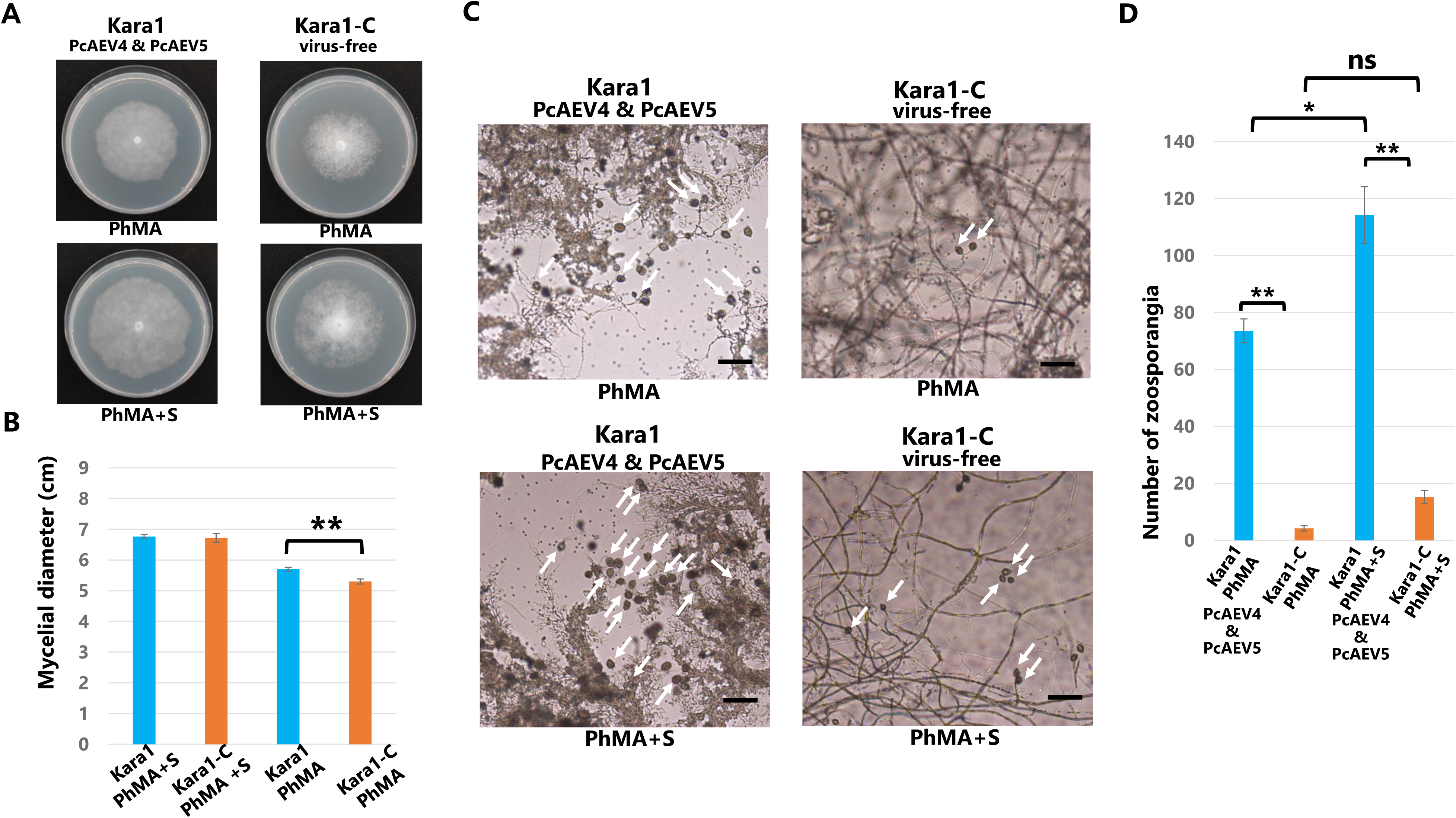
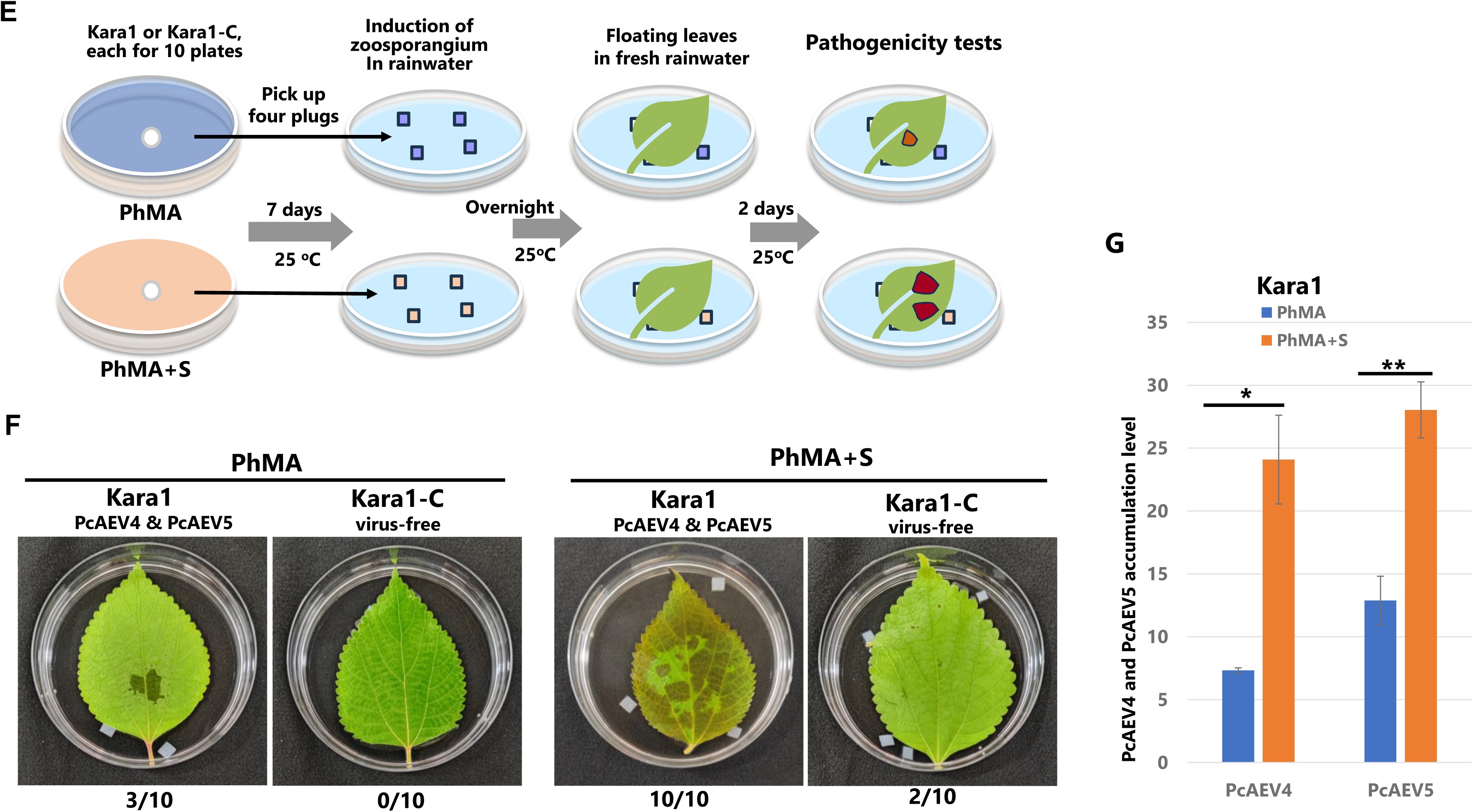
Effects of exogenous sterols on the Kara1 and Kara1-C strains. **(A)** Representative images showing morphologies and **(B)** histograms that compare mycelial diameters of the colonies of the Kara1 and Kara1-C strains after 7 days of culture on PhMA or PhMA+S (PhMA containing 25 μg/ml β-sitosterol) medium. **(C)** Representative images and **(D)** histogram comparing number of sporangia formed by the Kara1 and Kara1-C strains after 7 days of culture on PhMA or PhMA+S medium. Zoosporangia induction was performed by incubating hyphae cultured on cellophane membranes in DW under light conditions for 40 hours, and sporangia were counted in randomly selected fields. **(E)** Pathogenicity assay with sporangia on detached *B. nivea* var. *nipononivea* leaves induced by the Kara1 and Kara1-C strains. Diagram of the sporangia-induced pathogenicity test. **(F)** Lesions induced by the Kara1 and Kara1-C strains on *B. nivea* var. *nipononivea* leaves at 2 dpi. Ten detached leaves were used for the pathogenicity test, with representative results shown. **(G)** qRT-PCR analysis of PcAEV4 and PcAEV5 RNA levels in the Kara1 strain grown on PhMA and PhMA+S media. WS21 (40S ribosomal protein S3A) was used as an internal control. Error bars represent standard deviations from five biological replicates of a representative experiment. Statistical analysis was performed using Student’s t-test or Tukey-Kramer test (*P < 0.05, **P < 0.01). ns indicates no significant difference.

We next examined the effect of exogenous sterols on zoosporangia formation. The Kara1 strain demonstrated increased zoosporangia formation (approximately 1.6-fold more) (P<0.01) on PhMA+S medium than on PhMA medium (Fig. 5C, white arrows, Fig. 5D). The Kara1-C strain showed almost no zoosporangia formation on PhMA without sterols. Kara1-C demonstrated a tendency toward increased sporangia formation on PhMA+S medium as compared to PhMA medium, although this difference was not significant (Fig. 5C, white arrows, Fig. 5D). In addition, to examine the impact of sterols in the culture medium and the infection by PcAEV4 and PcAEV5 on fungal pathogenicity, we compared the pathogenicity of the Kara1 strain or the Kara1-C strain using their mycelium plugs as inoculation sources. The inoculum was prepared as shown in Fig. 5E, and the pathogenicity test was carried out on 10 leaves of *B. nivea* var. *nipononivea*. Kara1 grown on PhMA+S caused blight spots over the entire leaf in all inoculated leaves, while the disease incidence of Kara1 grown on PhMA and Kara1-C grown on PhMA+S was low at 20-30%, and the size of the blight spots were smaller (Fig. 5F). Kara1-C grown on PhMA did not show any blight symptoms at all (Fig. 5F). These results suggested that both co-infection with PcAEV4 and PcAEV5 and addition of exogenous sterols contribute to pathogenicity of *Phytophthora* spp.

We next examined the influence of exogenous sterols on the replication of PcAEV4 and PcAEV5. After Kara1 was cultured for 7 days on either PhMA or PhMA+S, total RNA was extracted and the levels of PcAEV4 and PcAEV5 RNA were quantified using RT-qPCR. The results revealed a significant increase in the accumulation of PcAEV4 and PcAEV5 RNA in cells grown on PhMA+S as compared to those grown on PhMA without sterols, with approximately 3.2-fold (P<0.05) and 2.2-fold (P<0.01) increases, respectively (Fig. 5G). These findings suggested that exogenous sterols enhance the replication of PcAEV4 and PcAEV5.

### 6. PcAEV4 and PcAEV5 localize to the ER membrane fraction

Since PcAEV4 and PcAEV5 RNA levels varied depending on the presence of β-sterols in the medium, it was hypothesized that membrane components may be involved in replication of these endornaviruses. To confirm this hypothesis, we partially purified several membrane fractions by cellular fractionation and examined which membrane fractions were enriched in dsRNA of the endornaviruses. Total nucleic acids were extracted from eight fractions (P1000, S1000, P15000, S15000, P20000, S20000, P100000, and S100000) and then subjected to agarose gel electrophoresis (Fig. 6A). As a result, dsRNA components that appeared to be derived from PcAEV4 and PcAEV5 were detected specifically in three pellet fractions: P15000, P20000, and P100000 (Fig. 6B). To further investigate whether PcAEV4 and PcAEV5 were associated with the mitochondrial fractions, P15000 (rich in intact mitochondria) and P20000 (rich in non-intact or small mitochondria) were separately subjected to ultracentrifugation in discontinuous sucrose density gradients (1.2 M, 1.3M, and 2.0M). In P15000, mitochondrial genomic DNA (Mito gDNA) was predominantly detected near the boundary between 1.3 and 2.0 M (fraction 11) (Fig. S2, lane 11), while PcAEV4 and PcAEV5 dsRNA were detected only in fractions 1 and 2 (Fig. S2, lanes 1 and 2). In P20000, Mito gDNA was detected in broad fractions, likely due to the precipitation of broken mitochondrial inner membrane, while PcAEV4 and PcAEV5 dsRNA was detected mainly in fractions 1 and 2, similar as in P15000 (Fig. S3, lanes 1 and 2). These results suggested that PcAEV4 and PcAEV5 are present in the lighter density fraction than the mitochondria. To test whether viral dsRNA in the membrane fraction is protected, P15000, P20000, and P100000 were treated with the nonionic surfactant, Triton-X, and RNase A. We found that only viral dsRNA in the P100000 membrane fraction showed RNase resistance, indicating that dsRNA in this fraction was protected by the membrane (Fig. 6C). To further purify the membrane fractions containing PcAEV4 and PcAEV5 dsRNA, the P100000 suspension was subjected to a continuous sucrose density gradient from 10% to 60%. Agarose gel electrophoresis of total nucleic acids extracted from the resultant 15 fractions revealed PcAEV4 and PcAEV5 dsRNA in fraction 7 (sucrose concentration, 25.6 %) (Fig. 6D, 6E). Northern blot analysis using strand-specific riboprobes specific for PcAEV4 and PcAEV5 revealed strong signals for the positive and negative strands of both endornaviruses in fraction 7 (Fig. 6G-6J, lanes 6-9). A nick specific to the plus strand was also detected (Fig. 6G, 6I, lanes 6-9).

**Figure 6.**
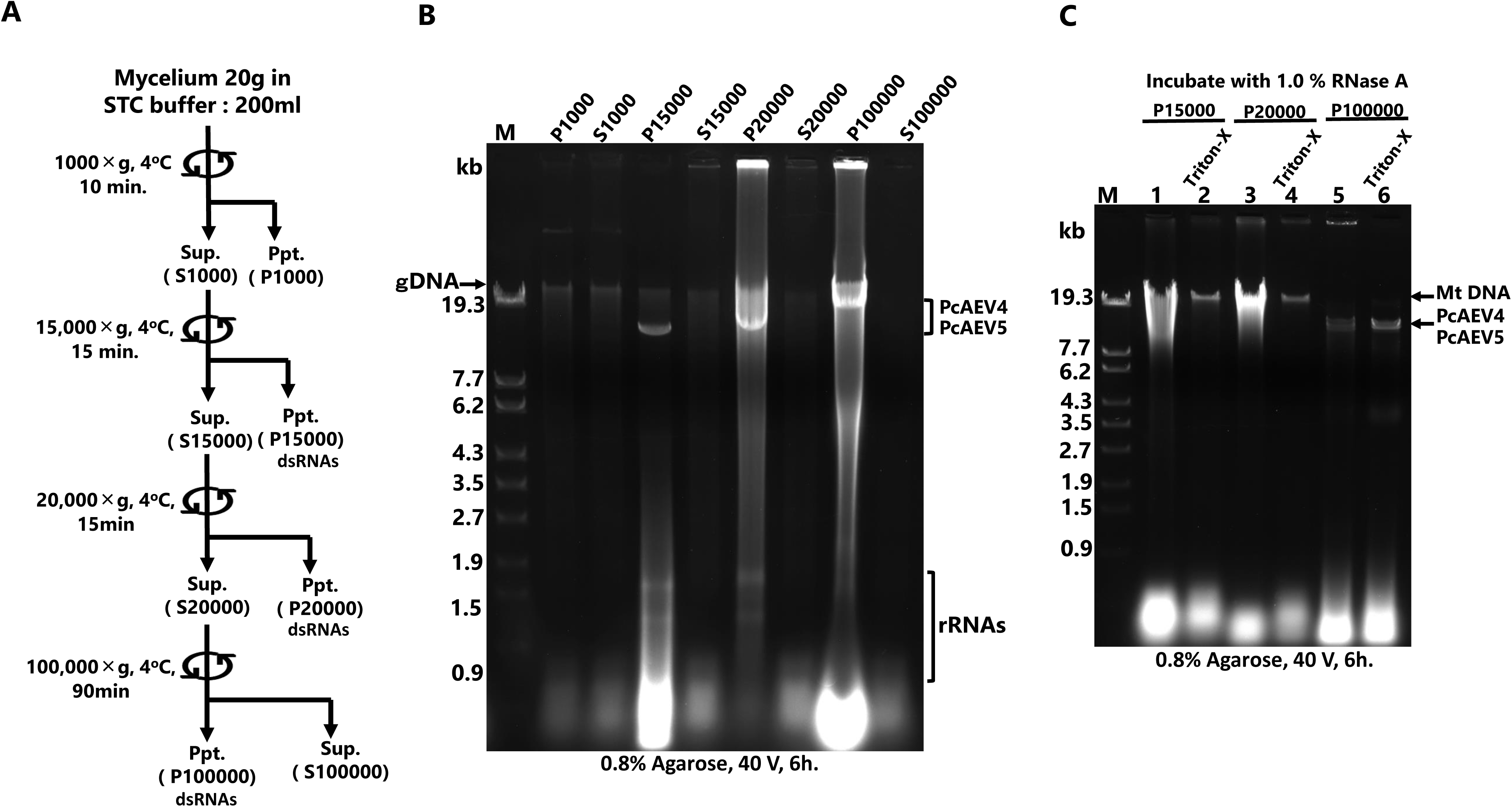
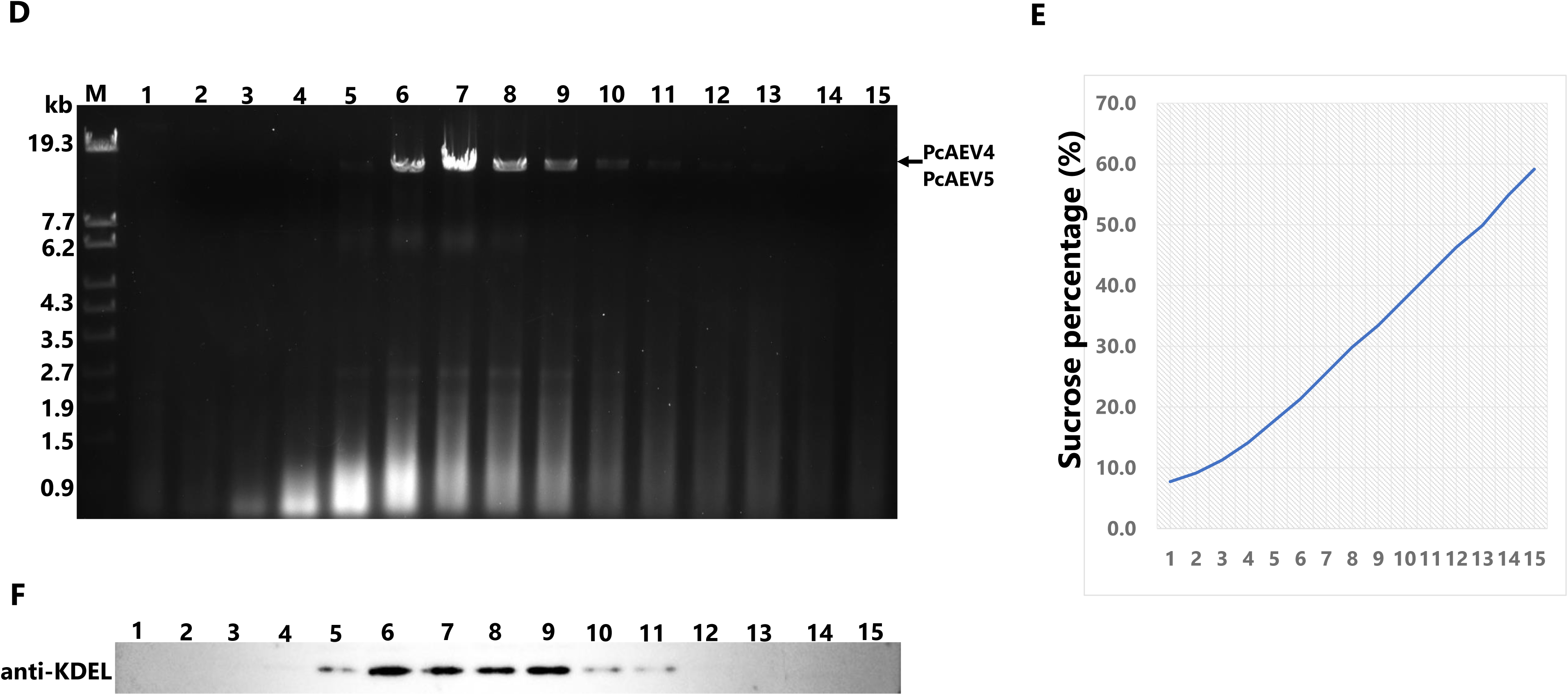
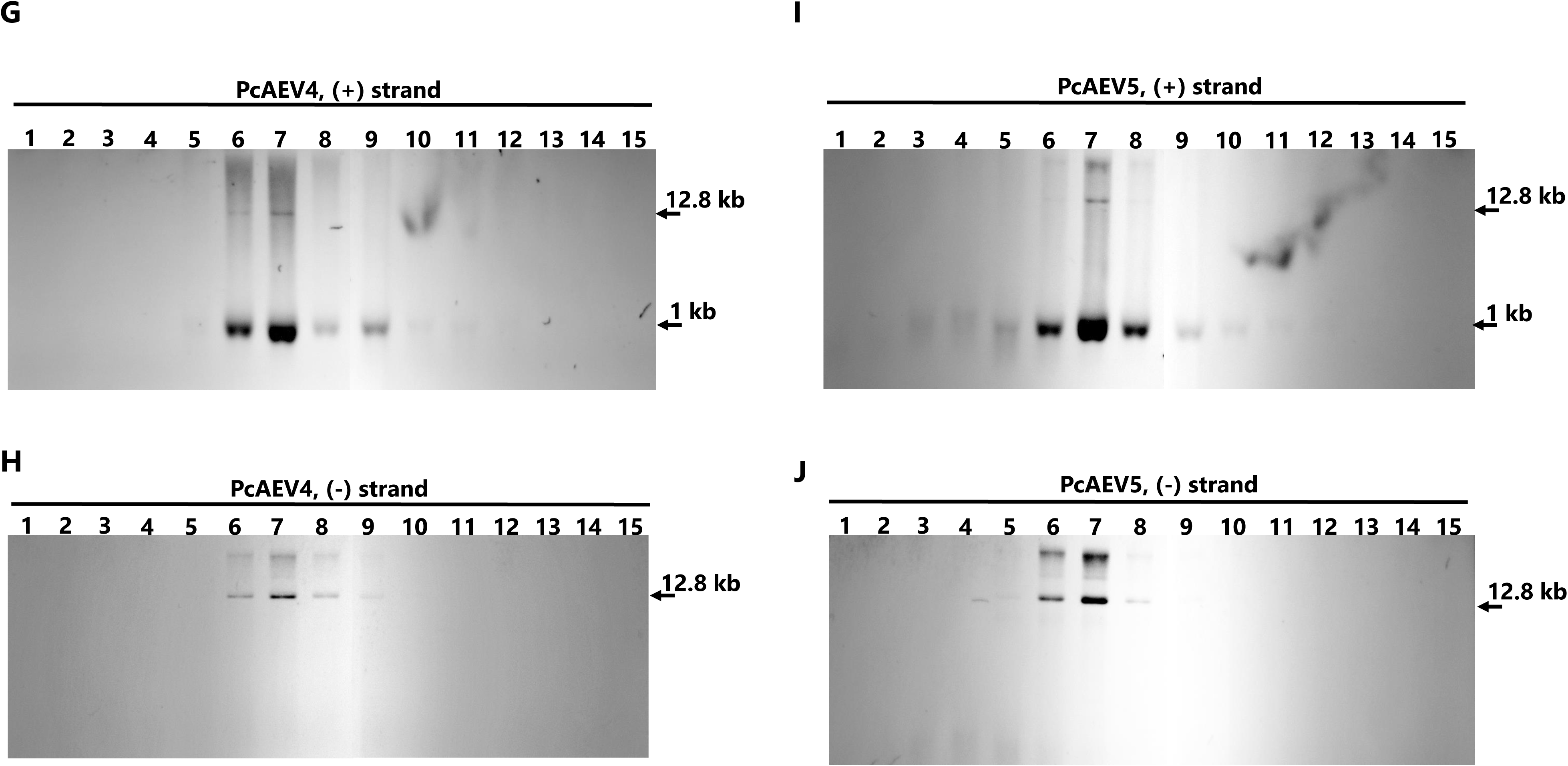
Replication intermediates of PcAEV4 and PcAEV5 are concentrated in ER membrane fractions. **(A)** Overview of membrane fraction purification in the Kara1 strain. Crushed cell lysates were centrifuged at 1,000×g, 15,000×g, 20,000×g, and 100,000×g. Pellets (Ppt.) and supernatants (Sup.) were collected and analyzed. **(B)** Electrophoresis of total nucleic acids extracted from each fraction. Electrophoresis was performed on 0.8% agarose gel at 80V for 2 hours, followed by staining with ethidium bromide (0.5 μg/ml). Note the different levels of concentration between pellet (P) and supernatant (S). Lane designations: M: DNA marker (250 ng of λDNA digested with EcoT14I). The arrow indicates the position of the 12.8 kb dsRNA. **(C)** RNase A sensitivity assay of PcAEV4 and PcAEV5. Portions of membrane fractions obtained by cell fractionation were incubated in isotonic solution containing RNase A at 37°C for 30 minutes. Electrophoresis was performed using P15000, P20000, and P100000 fractions with (right lanes) or without (left lanes) 1% Triton X-100. Lane designations: M: DNA marker (250 ng of λDNA digested with EcoT14I). The arrows indicate the positions of mitochondrial genomic DNA (Mt DNA) and PcAEV4 and PcAEV5 dsRNA. **(D)** Sucrose gradient fractionation of microsomal fraction P100000. Fractions obtained were labeled 1 through 15 from top to bottom (lanes 1-15). Total nucleic acids were extracted from 1/10 of each fraction and subjected to agarose gel electrophoresis. Lane M: DNA marker (250 ng of λDNA digested with EcoT14I). The arrow indicates the position of PcAEV4 and PcAEV5 dsRNA. **(E)** The percentage of sucrose concentration was determined by refractometry. **(F)** Western blot assay with anti-KDEL antibody was performed after concentrating the obtained fractions tenfold, with equal amounts of protein loaded in all lanes. **(G, H, I, J)** Northern blot analysis of the sucrose gradient fractions of microsomal fraction P100000. Approximately 20 µg of total nucleic acids extracted from the obtained fractions was loaded, and DIG-labeled RNA probes were used for detection. Probes were prepared at the following positions: PcAEV4-5’ (nt 114-837) **(G)**, PcAEV4-3’ (nt 11,610-12,037) **(H)**, PcAEV5-5’ (nt 101-791) **(I)**, and PcAEV5-3’ (nt 11,547-12,331) **(J)** probes. Arrows indicate the positions of PcAEV4 and PcAEV5 genomic RNA (12.8 kb) and nicks (1 kb).

To investigate whether PcAEV4 and PcAEV5 are associated with the ER, we performed immunoblotting analysis for KDEL, an ER marker, and found that KDEL signals peaked in fractions 6, 7, 8, and 9, and their localizations were consistent with PcAEV4 and PcAEV5 (Fig. 6F). These results suggested that the dsRNA replication intermediate of these endornaviruses localize to the ER membrane.

### 7. Visualization of PcAEV4 and PcAEV5 dsRNA by immunofluorescence assay (IFA)

In order to visualize the localization of PcAEV4 and PcAEV5 in host cells, we performed IFA using the J2 antibody, which binds specifically to dsRNA molecules [39]. As a negative control, protoplasts were also prepared from the virus-cured strain Kara1-C, and differences in labeling intensity of dsRNA detected in the cells were compared (Fig. S4).

As shown in Fig. 7A, numerous small granule-like structures (green) corresponding to J2 antibody-targeted dsRNA molecules were observed in the protoplasts of the PcAEV4- and PcAEV5-infected Kara1 strain. On the other hand, very few if any granule-like structures were observed in the Kara1-C strain, which was expected as this strain was cured of the endornaviruses (Fig. 7A, right panels).

**Figure 7.**
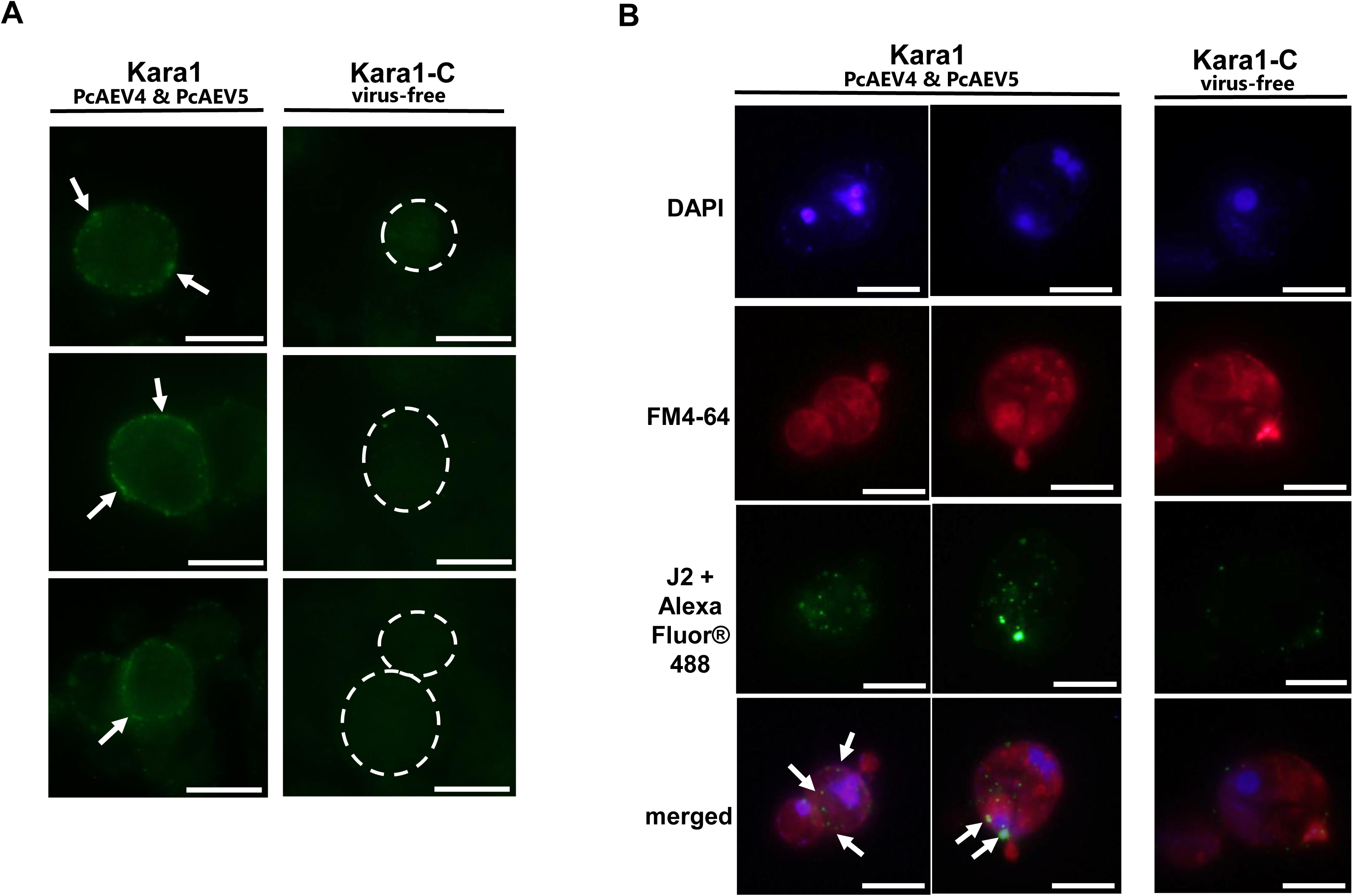
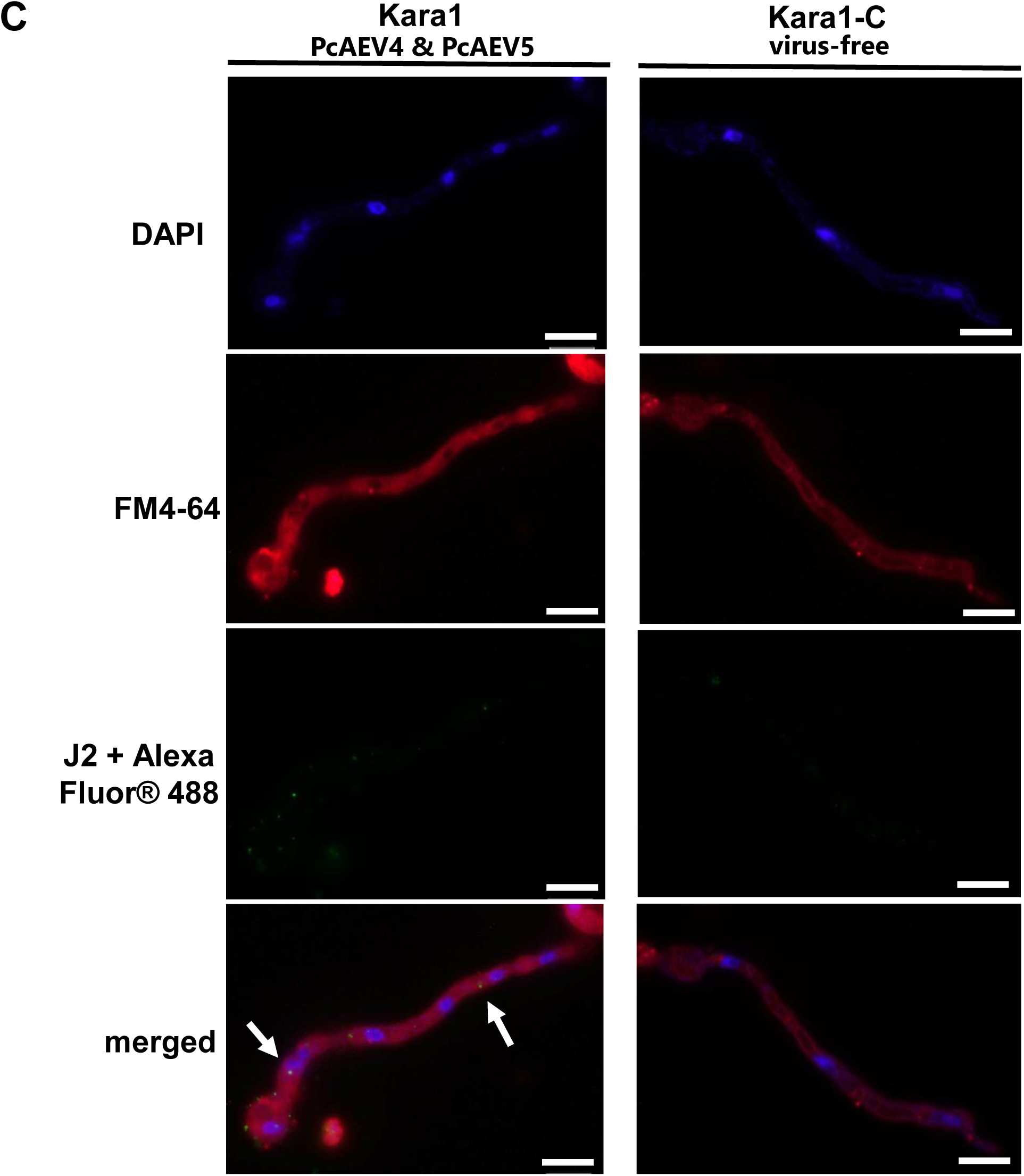
Visualization of endornavirus-derived dsRNA co-localized with membrane components by the dsRNA-specific antibody J2. **(A)** Detection of endornavirus-derived dsRNA labeled with J2 antibody in protoplasts of the Kara1 (left panels) and Kara1-C (right panels) strains. White arrows indicate representative green fluorescence strongly stained by J2 antibody, and white dashed lines outline cell positions. **(B) (C)** Co-localization of endornavirus-derived dsRNA with lipid membranes stained by FM4-64 in regenerated protoplasts or regenerated mycelium of the Kara1 (left and middle panels) and Kara1-C (right panels) strains. **(B),** protoplasts were regenerated overnight in regeneration medium, followed by lipid membrane staining with FM4-64, and then fixation. DAPI staining was performed after the primary and secondary antibody reactions. White arrows indicate representative co-localization signals of lipid membranes stained with FM4-64 and dsRNA stained with J2 antibody. Kara1-C strain was used as a negative control. **(C),** regenerated mycelium. Scale bar = 10 μm.

Next, to investigate the association of dsRNA with lipid membranes, IFA was performed on regenerated protoplasts and mycelium stained with FM4-64. In the regenerated protoplasts, the intensity of fluorescence was higher than that observed in the protoplasts (Fig. 7B), suggesting a high accumulation of PcAEV4 and PcAEV5 dsRNA in the cytoplasm (Fig. 7B, red staining). Co-staining with the lipid membrane dye FM4-64 revealed partial co-localization of J2 antibody-labeled granule-like structures with FM4-64-labeled aggregates. (Fig. 7B, green staining, white arrows). Granule-like structures were also observed in the regenerated mycelium, which also suggested that endornavirus dsRNA accumulated in the cytoplasm, although they showed weaker fluorescence than those in the regenerated protoplasts (Fig. 7C). The granule-like structures in the regenerated mycelium showed weaker fluorescence than those in the regenerated protoplasts, which may have been due to insufficient penetration of the antibody and/or fluorochrome through the cell wall of the mycelial cells. These results indicated that endornavirus dsRNA associates with cytoplasmic lipid membrane components. Note that a few granule-like structures labeled with J2 antibody were also observed in the endornavirus-free Kara1-C strain (Fig. 7B, right panel), which were assumed to be derived from highly structured ribosomal RNA in the cytoplasm and not endornavirus dsRNA.

## Discussion

In recent years, reports of severe damage caused by the forest pathogen *Phytophthora* spp. have been increasing, and attention has focused on *Phytophthora* spp. isolated from the natural environment of forests and rivers. The search for viruses in *Phytophthora* spp. has expanded to include those living in forests in addition to agricultural fields [10]. In this study, two novel endornaviruses were identified in *P. cactorum* isolated from a forest ecosystem in Japan. The total number of alphaendornaviruses infecting *Phytophthora* spp. reported to date is six, including PcAEV4 and PcAEV5 reported here, PcAEV1, PcAEV2, and PcAEV3 isolated from the Finnish strawberry blight pathogen *P. cactorum* (Supplemental Fig. S1) [8], and Phytophthora cactaceae RNA virus 5, presumed to be an endornavirus that infects a different species (*P. cactaceae)*. Interestingly, PcAEV5 in this report was most closely related to PcAEV3 from Finland. This suggests that the virus-infected *P. cactorum* strain found in the natural ecosystem of Japan may have become pathogenic to strawberries across continents or spread among the same *P. cactorum*, and that the alphaendornaviruses may be key RNA molecules in inferring the movement of plant pathogenic *Phytophthora* spp.

Since *Phytophthora* spp. do not have a septum, it is assumed that endornaviruses in this fungus can be distributed throughout the mycelium, making it difficult for us to eliminate the virus. Further, the vertical transmission rate of PEV2 and PEV3 after single zoospore isolation was as high as 100% [9]. We therefore devised a combined treatment of protoplastization and antiviral agents (ribavirin and cycloheximide) for the Kara1 strain, which was co-infected with PcAEV4 and PcAEV5, and successfully isolated at least 16 virus-free strains (Fig. 3. Fig. S4A). In order to reduce the potential impact of genomic mutations that may have been incurred during the process of mycovirus curing using these agents [40], experiments were conducted with 5 biological replicates.

The PcAEV4- and PcAEV5-free Kara1-C strain showed increased mycelial growth as compared to the original Kara1 strain, but significant reduction in zoosporangia formation (Fig. 3C-3F), as well as reduced lesion formation on the leaves of *B. nivea* var. *nipononivea* (Fig. 3G-3H), indicating reduced virulence. Despite the enhanced growth, the Kara1-C strain showed reduced resistance to high temperature (Fig. 4A, 4B), high osmotic pressure, cell wall synthesis inhibition (Fig. 4C, 4D), and cell wall digestion (Fig. 4E). These results suggested that virus curing may result in weakening of the cell wall, leading to decreased virulence and reduced resistance to various stresses. The endornavirus-free Kara1-C strain was also more susceptible to Calcofluor white, which inhibits cellulose synthesis, than to Congo red, which inhibits beta-glucan polymerization (Fig. 4C, 4D), suggesting that PcAEV4 and PcAEV5 co-infection may induce cellulose biosynthesis in mycelia. A report that knockout of the cellulose synthase gene PcCesA1, which is highly conserved among *Phytophthora* spp., in *Phytophthora capsici* had negative effects on mycelial growth, germination of cystospores, and virulence [41] supports our findings here. There are a few reports on the involvement of virus infection in cell wall biosynthesis; in *Penicillium stoloniferum*, *Aspergillus foetidus*, and *Aspergillus niger*, virus-infected strains produced more galactosamines than uninfected strains, and in *Botrytis cinerea,* the viruses induced production of chitin/glycoproteins that were included in the cell wall [42, 43, 44].

Viral infections were suggested to promote zoosporangia formation in PEV2- and PEV3-infected *Phytophthora* pathogen of asparagus and PiRV2-infected *P. infestans* [12, 9], which corroborated our results in PcAEV4- and PcAEV5-infected *P. cactorum*. Poimala et al. found that infection with Phytophthora cactorum bunya-like virus 1 and 2 in the strawberry blight fungus *P. cactorum* significantly reduced host mycelial growth, sporangium production, and size, but did not alter pathogenicity [45]. Considering these results, the effects on growth and virulence of plant pathogens by viruses appear to differ depending on the type of mycovirus infecting the host oomycete, and that the growth situation in culture medium is not correlated with virulence.

Experimental introduction of viruses is important to determine the effects of viral infections. However, in this study, we were unable to successfully propagate PcAEV4 and PcAEV5 by confrontation culture (data not shown). In fact, to date, virus transmission via the confrontation culture method in *Phytophthora* has only been reported for PiRV2 and has not yet been successful for PiRV3, PcBV1, and PcBV2 [6, 12, 45]. Since *P. cactorum* is a homothallic species (self-fertile), a strong barrier line is observed in confrontation cultures between homologous strains, which is not appropriate for mycelial fusion experiments [46]. In the future, using heterothallic species of *Phytophthora* (types A1 and A2) in the confrontation culture method could lead to successful propagation of these viruses.

Despite the lack of a sterol biosynthetic pathway, *Phytophthora* spp. can grow *in vitro* on sterol-free medium, whereas exogenous addition of sterols promotes mycelial growth and reproduction [32, 47]. In this study, the addition of sterols to the medium also promoted mycelial growth (Fig. 5A, 5B), zoosporangia formation (Fig. 5C, 5D), and enhanced virulence in the PcAEV4- and PcAEV5-infected Kara1 strain (Fig. 5F). Interestingly, sterols significantly increased the accumulation of PcAEV4 and PcAEV5 RNA in the mycelium (Fig. 5G). Recently, the PcDHCR7 gene, which converts ergosterol to brassicasterol, was discovered in *Phytophthora capsici* [48]. Further, pathways related to sterol biosynthesis, such as the mevalonic acid pathway, and modification of exogenous sterols as well as four genes with sterol-sensing domains (SSD) were discovered in *Phytophthora sojae* [49, 50]. These findings suggested that a sterol-induced intracellular signaling network exists in the genus *Phytophthora*. Interestingly, the PcAEV4- and PcAEV5-infected Kara1 strain maintained hyphal elongation, zoosporangia formation, and replication of endornaviruses even in sterol-depleted medium (Fig. 5C, 5D), suggesting that PcAEV4 and PcAEV5 co-infection positively affects host sterol metabolism and exogenous sterol-sensing mechanisms, which enhance virulence-associated traits in the host.

In previous reports on the plant endornaviruses, Vicia faba endornavirus 1 [18] and Oryza sativa endornavirus 1 [51, 52], their RdRp activities were detected in host membranous vesicles, while Bell pepper endornavirus 1 infection caused slight morphological changes in host mitochondria and chloroplasts [53], suggesting that plant endornaviruses are associated with intracellular membrane components. In this study, we demonstrated that endornavirus-specific dsRNA, a replication intermediate, co-localized with KDEL, suggesting an ER subcellular localization (Fig. 6F). Many viruses in the order *Martellivirales* to which endornaviruses belong, such as Tobacco mosaic virus and Bromovirus, were reported to replicate by remodeling membrane components derived from the ER [21, 22]. In support of this, we detected dsRNA aggregates (shown in green) inside the plasma membrane in protoplasts of the Kara 1 strain (Fig. 7A), and then larger green aggregates were detected in regenerated protoplasts after 20 hours at 25°C (Fig. 7B). These results suggested that the dsRNA replication intermediates of PcAEV4 and PcAEV5 are associated with the ER membrane, which could protect against RNase A (Fig. 6C). Although the localization of dsRNA detected in protoplasts and regenerating protoplasts differed, which may be because protoplast regeneration may be accompanied by cell wall regeneration and large-scale modification of the intracellular membrane system that in turn changes the localization of the VRC.

In the PcAEV4- and PcAEV5-infected Kara1 strain, FM4-64 staining revealed many lipid drop (LD)-like structures in the mycelium (Fig. 7C), which may be involved in the putative VRC. LDs are cell organelles composed of a monolayer of phospholipid membranes derived from the ER and a neutral lipid core containing sterols, and are usually circular in shape with a size of 100 nm to 100 μm [54]. It is possible that infection with these endornaviruses promotes LD formation in mycelial cells; viruses that use LD for replication, such as Dengue virus, Hepatitis C virus, and rotaviruses, have been reported among animal viruses [55, 56, 57]. The localization of these two endornaviruses’ dsRNA replication intermediates in the ER membrane suggested that they modulate lipid metabolism in the host *P. cactorum*, which can enhance its virulence. In the future, we would like to comprehensively investigate the effects of the endornaviruses infection by transcriptomic and lipidomic analyses.

## Materials and Methods

### Fungal Strains and Culture Conditions

*P. cactorum* strain Kara1 was isolated from *B. nivea* var. *nipononivea* exhibiting wilt symptoms in Tochigi Prefecture, Japan. All strains, including the virus-cured strain Kara1-C, were cultured in darkness for 14 days and then maintained on modified Weitzman-Silva-Hunter agar medium (1.0% oatmeal, 0.1% Mg_2_SO_4_, 0.1% 7H_2_0. KH_2_PO_4_, and 0.1% NaNO_3_). For all experiments involving strain propagation, mycelia were pre-cultured on Phytophthora minimal agar (PhMA) medium (0.1 g KNO_3_, 0.2 g K_2_HPO_4_, 0.1 g MgSO_4_, 0.1 g CaCl_2_, 0.1 g L-asparagine, 0.05 g L-serine, 4 g glucose, and 1 mL trace elements [200 mg FeEDTA, 10 mg CuSO_4_, 10 mg MnCl_2_, 10 mg Na_2_MoO_4_, 10 mg Na_2_B4O_7_, 20 mg ZnSO_4_, and 100 mg thiamine hydrochloride in 100 mL of distilled water] in 1 L of distilled water) at 25°C in darkness for 7 days [50, 58].

### Total RNA Extraction, dsRNA Purification, and RT-PCR

Total RNA extraction was performed using mycelia grown in liquid medium at 25℃. According to the manufacturer’s instructions, total RNA was extracted from 0.05 g of mycelia using the RNeasy Plant Mini Kit (QIAGEN, Hilden, Germany). For dsRNA purification, spin columns containing cellulose D (Advantec, Tokyo, Japan) were used [64]. Mycelia (0.05 g dry weight) were ground in 500 µL of extraction buffer (100 mM NaCl, 10 mM Tris-HCl pH 8.0, 1 mM EDTA, 1% SDS, and 0.1% (v/v) β-mercaptoethanol) and mixed with an equal volume of phenol-chloroform-isoamyl alcohol (25:24:1). The aqueous phase containing total nucleic acids was mixed with ethanol (final concentration 16%), and dsRNA was selectively purified using the spin column method. Finally, dsRNA was precipitated with ethanol and stored at -80°C. Total RNA and dsRNA were quantified by agarose gel electrophoresis.

RT-PCR for sequencing the complete genomes of PcAEV4 and PcAEV5 and for preparing probes for Northern blot analysis was performed as follows; reverse transcription was carried out using total RNA or purified dsRNA extracted from strain Kara1 or Kara1-C as template and SuperScript® IV Reverse Transcriptase (Thermo Fisher Scientific, Waltham, MA, USA) according to the manufacturer’s instructions. PCR reactions were performed using the cDNA as a template with primers listed in Table S1 and GoTaq (Promega, Madison, WI, USA).

### Next-Generation Sequencing and Full-Length Sequence Determination

The PrimeScript^TM^ II 1st strand cDNA Synthesis kit (Takara Bio USA, San Jose, CA, USA) was used to prepare the sequencing library from dsRNA extracted from the Kara1 strain, and 8,571,034 reads were generated by next-generation sequencing using Illumina NovaSeq 6000 (San Diego, CA, USA). Sequence reads were assembled *de novo* using CLC Genomics Workbench version 11 (CLC Bio-QIAGEN, Aarhus, Denmark). The 371 assembled sequence contigs were subjected to virus sequence screening using local Basic Local Alignment Search Tool (BLAST) with virus reference sequences from the National Center for Biotechnology Information (NCBI, https://ncbi.nlm.nih.gov/).

Rapid amplification of cDNA ends (RACE) was used to determine 5’ and 3’ terminal nucleotide sequences of PcAEV4 and PcAEV5 with the SMARTer® RACE 5’/3’ Kit (Takara Bio USA) according to the manufacturer’s instructions. To amplify the 3’ end, total RNA extracted and purified from the Kara1 strain was polyadenylated by poly(A) tailing reaction before use. RACE-PCR products were TA-cloned into pGEM-T easy (Promega), and the nucleotide sequences were determined by the Sanger method. The primer pairs used are shown in Supplementary Table S1.

### Phylogenetic Analysis

The obtained nucleotide sequences were analyzed for ORFs and translated into amino acid sequences, then subjected to protein similarity searches using GENETYX software ver.9 (GENETYX, Tokyo, Japan). Multiple alignments based on estimated amino acid sequences were constructed by CLUSTAL_X ver 2.0 [59, 60] and MEGAX [61]. Phylogenetic analysis under the maximum likelihood method was performed using the optimal model of amino acid substitution selected by ProtTest 2.4 and PhyML 3.1 [65], with bootstrap tests performed with 1,000 resamplings.

### Northern Blot Analysis

Northern blot analysis was performed with slight modifications to the method of Sakuta et al., 2023 [13]. Specifically, approximately 10 µg of total RNA extracted from strain Kara1 was heat-denatured at 65°C for 5 min, then separated on a 0.8% agarose 3-(N-morpholino) propane sulfonic acid (MOPS) gel containing 6% formaldehyde, and transferred to a positively charged nylon membrane (Roche, Basel, Switzerland) by capillary blotting. To detect PcAEV4 and PcAEV5 genomes, DIG-labeled DNA probes for PcAEV4-5’ (nt 114-837), PcAEV4-3’ (nt 11,610-12,37), PcAEV5-5’ (nt 101-791), and PcAEV5-3’ (nt 11,547-12,331) were prepared with specific primer pairs (Table S1) using PCR DIG Labeling Mix (Roche). Riboprobes were also prepared using the DIG RNA Labeling Kit (SP6/T7) (Roche) according to the manufacturer’s protocol. DIG-labeled riboprobes (RNA probes) were obtained by runoff transcription using the same regions as the PcAEV4-5’ (nt 114-837) and PcAEV5-5 (nt 101-791) probes as templates. Hybridization using DNA and RNA probes was performed at 50°C and 65℃, respectively, for 16 h. After hybridization, the membrane was washed twice in low stringency buffer (2×SSC, 0.1% SDS) at 37°C for 5 minutes and twice in high stringency buffer (0.1% SSC, 0.1% SDS) at 50℃ for 15 minutes. Hybridization signals were detected with addition of ready-to-use CDP-Star® (Roche) and captured with Ez-Capture MG (ATTO, Tokyo, Japan).

### Protoplast Preparation and Isolation of Virus-Cured Strains

For protoplast preparation, mycelial plugs of strain Kara1 were inoculated into four 100 ml flasks each containing 25 ml of 10% V8 liquid medium and cultured at 25°C for 48 h without shaking. The cultured mycelia were collected on sterilized Miracloth (Merck, Darmstadt, Germany) and washed twice with distilled water. Subsequently, mycelia rinsed with 0.8 M mannitol were plasmolyzed in 0.8 M mannitol for 10 min. These mycelia were again filtered through sterilized Miracloth into 25 ml of enzyme solution for cell wall digestion (0.75% lysing enzyme [Merck] and 0.6% cellulase [Yakult, Tokyo, Japan]), centrifuged at 600×g for 3 min at 4°C to pellet the protoplasts, resuspended in 10 ml of W5 solution (2 mM 2-(N-morpholino) ethane sulfonic acid, 154 mM NaCl, 125 mM CaCl2, 5 mM KCl; pH 5.7), and pelleted again to wash the protoplasts. After confirming the state of protoplasts under an optical microscope, they were resuspended in W5 solution to 1×10⁶/ml to prepare the protoplast preparation.

Ribavirin and cycloheximide (Fujifilm Wako, Osaka, Japan) were added to the protoplast suspension to final concentrations of 300 µg/mL and 5 µg/mL, respectively. Additionally, Regeneration V8, containing 0.4 M mannitol and 5% V8, was added at twice the volume of the protoplast suspension. The mixture was incubated at 25°C in the dark for 20 h to regenerate the cell walls of the protoplasts. Regenerated mycelia were inoculated onto Regeneration V8A (A: agar) medium containing ribavirin (300 µg/ml), cycloheximide (5 µg/ml), and ampicillin (50 µg/ml), and incubated at 25°C until colonies appeared. Sixteen single colonies were transferred to V8A medium containing ribavirin and cycloheximide by the hyphal tip isolation. This procedure was repeated four additional times (a total of 5 times on V8A medium with drug treatment), after which the isolates were subcultured on V8A medium without drugs. The subcultured mycelia were cultured in V8 liquid medium and curing of virus was confirmed by dsRNA purification and RT-PCR.

### Analysis of Pathogenicity and Zoosporangium Formation

Pathogenicity of Kara1 and Kara1-C strains was compared using leaves of wild-growing *B. nivea* var. *nipononivea*. Collected leaves were washed under running water for 1-2 h before use. In the test shown in Fig. 3G, mycelial plugs of both strains cultured on V8A medium for one week, and V8A plugs as a control, were placed on the same intact leaf, and the lesion areas formed after 4 dpi at 20°C were compared by calculating the area using ImageJ [62]. In the tests shown in Fig. 5E and 5F, mycelial plugs of strains Kara1 or Kara1-C cultured for one week on PhMA or PhMA+S containing 25 µg/ml β-sitosterol were incubated overnight in Petri dishes filled with distilled rainwater to promote zoosporangium formation. The next day, the water was replaced with fresh rainwater, and washed leaves of *B. nivea* var. *nipononivea* were floated on the water surface, and the infection rate at 20°C for 2 dpi was compared.

For comparison of the number of zoosporangium, sterile cellophane membranes (1.0 cm x 1.0 cm) were placed on V8A, PhMA, or PhMA+S medium. After 10 days of inoculation when the mycelia completely covered the plate, the cellophane membrane was carefully peeled off with tweezers, submerged in a Petri dish filled with rainwater, and incubated at 20°C overnight to promote zoosporangium formation. The next day, the number of zoosporangia was counted five times to calculate the average value.

### RT-qPCR

RT-qPCR was performed using total RNA as a template after removing contaminating genomic DNA using gDNA Eraser (TaKaRa, Shiga, Japan). The reaction was carried out using 100 ng of total RNA with the Thermal Cycler Dice Real-Time System and GoTaq® 1-step RT-qPCR Systems (Promega). The accumulation of PcAEV4 and PcAEV5 RNA in Kara1 was relatively quantified by normalizing to the reference gene, PcWS21, using the primer set, PcWS21_F and PcWS21_R (Yan et al., 2006). The qPCR primers, PcAEV4-qrt_F, PcAEV4-qrt_R, PcAEV5-qrt_F, and PcAEV5-qrt_R, were designed using the Primer3Plus program (http://www.bioinformatics.nl/cgi-bin/primer3plus/primer3plus.cgi/) from Sol Genomics Network (https://solgenomics.net/). RT-qPCR primer sequences are listed in Table S1.

### Collection of Membrane Fractions by Centrifugal Fractionation

Approximately 20 g of mycelia were homogenized in 200 ml of isotonic solution (0.25 M sucrose, 100 mM Tris-HCl, pH 7.4, 10 mM EGTA). This homogenate was transferred to a 50 ml centrifuge tube and centrifuged at 1000×g for 10 min to precipitate cell debris and nuclei (P1000). The supernatant was transferred to Hyzex tubes and centrifuged at 15000×g for 15 min to obtain a fraction rich in mitochondria (P15000). This supernatant was transferred to another Hyzex tube and centrifuged at 20000×g for 15 min, which was considered the membrane fraction (P20000). Furthermore, this supernatant was centrifuged at 100000×g for 90 min using a P45AT rotor in a Hitachi CP80WX (Hitachi, Koki, Japan), and the precipitate was considered the microsomal fraction (P100000) and the supernatant was the cytosol fraction (S100000). The obtained fractions were mixed with 0.5 ml of 2×STE buffer (200 mM NaCl, 20 mM Tris-HCl pH 8.0, 2 mM EDTA pH 8.0) containing 1% SDS and 0.5 ml of 1:1 phenol/chloroform. The mixture was vortexed and centrifuged at 15,000×g for 5 min. The supernatant was used as the total nucleic acid sample, and the presence of endornaviruses was confirmed by agarose gel electrophoresis.

Next, P15000 and P20000 obtained above were subjected to discontinuous sucrose density gradient centrifugation. Sucrose solutions of 1.2 M, 1.3 M, and 2.0 M were prepared in TE buffer (10 mM Tris-HCl pH 7.4, 1mM EDTA) and gently layered. Centrifugation was performed at 82200×g for 3 h using a P28S swing rotor in a Hitachi CP80WX, and divided into 14 fractions of 2 ml each. Total nucleic acids were extracted from 1/10 of the obtained fractions by the same method as described above and subjected to agarose gel electrophoresis.

For P100000, to obtain more detailed fractions, it was subjected to 10 - 60% sucrose density gradient centrifugation. Centrifugation was performed at 82200×g for 3 h using a P28S swing rotor in a Hitachi CP80WX, and 15 fractions of 2 ml each were obtained. To concentrate the obtained fractions, centrifugation was performed at 100000×g for 1.5 h using a P70AT rotor in a Hitachi CP80WX, and the precipitate was suspended in 100 µl of isotonic solution to obtain concentrated membrane fractions.

### RNase A Sensitivity Test for Membrane Fractions

RNase A digestion of the samples was performed together with or without 1% (v/v) Triton-X at 37°C for 30 min. After RNase A digestion, samples were extracted with phenol-chloroform and precipitated with isopropanol. Following a 70% EtOH wash and drying step, each sample was dissolved in 10 µl of distilled water. The presence of viral dsRNA was confirmed by electrophoresis.

### Gel Electrophoresis and Immunoblotting

Concentrated membrane fractions were separated by 8% SDS-PAGE in 25 mM Tris-glycine and 0.1% SDS at 20 mA for 1.5 h. Proteins were electrotransferred to PVDF membranes using the AE-6677 semi-dry electrophoretic transfer system (ATTO). Membranes were treated with blocking buffer (5% (w/v) skim milk, 0.2% (v/v) Tween 20 in PBS). Anti-KDEL mouse monoclonal antibody (1:1,000, MEDICAL & BIOLOGICAL LABORATORIES, Tokyo, Japan) and secondary goat anti-mouse antibody-horseradish peroxidase conjugate (1:10,000, MEDICAL & BIOLOGICAL LABORATORIES) were used according to the manufacturer’s instructions. Signals were detected with ECL Plus Western Blotting Detection Reagent (GE Healthcare, Chicago, IL, USA) and visualized using the Ez-Capture MG imaging system (ATTO).

### Immunofluorescence Assay (IFA)

IFA was performed following the previously described method [66] with slight modifications. Prepared protoplasts were fixed with 4% paraformaldehyde (PFA) at room temperature for at least 30 min. Autofluorescence quenching was performed in PBS containing 0.1 M glycine for 10 min. Cell permeabilization was carried out with PBS containing 0.1% saponin for 10 min, followed by blocking with PBS containing 5% fetal calf serum (FCS) for at least 1 h. Immunostaining was performed as follows: for primary antibody reaction, dsRNA-specific J2 antibody (1:250, Nordic-MUbio BV, Susteren, Netherlands) was added and incubated at room temperature for at least 1 h. After reaction, the antibody solution was discarded, washed three times with PBS, and the secondary antibody reaction was performed with anti-mouse IgG conjugated with Alexa 488 (1:1.000, Thermo Fisher Scientific) for 1 h at room temperature. After cells were washed three times with PBS, they were suspended in glycerol and spread on a slide glass, sealed with a cover glass and nail polish. Fluorescence microscopy observations were performed using an optical fluorescence microscope (Olympus IX71, Tokyo, Japan).

IFA on regenerated protoplasts was performed as follows: protoplasts were regenerated overnight in Regeneration V8 liquid medium, washed with PBS, and stained with 5 µg/mL FM4-64 (Thermo Fisher Scientific) for 10 min in the dark. After staining, fixation was performed with 4% PFA at room temperature for at least 30 min, followed by immunostaining as described above. After immunostaining, 1 µg/ml 4’,6-diamidino-2-phenylindole (DAPI, Merck) was added for nuclear staining and incubated at room temperature for 10 min. After washing with PBS, samples were mounted and observed under a fluorescence microscope.

### Statistical Analysis

Colony diameter measurement data were subjected to analysis of variance. Significant differences between control and treatment groups were analyzed using one-way analysis of variance (ANOVA), and statistical significance was analyzed using Student’s *t*-test. Pathogenicity comparison between Kara1 and Kara1-C strains was quantified by measuring the formed lesion areas using ImageJ [62], following the instructions in the ImageJ manual. For zoosporangium count measurements, differences between means of different treatments were analyzed using the Tukey–Kramer test. Data are presented as mean ± standard deviation (SD). Statistical significance was set at *p* < 0.05 (* *p* < 0.05, ** *p* < 0.01). The sample size (n) for each statistical analysis is shown in the corresponding figure legend.

## Supporting information

Supplemental Data 1-4, and Certificate

## Author contributions

KS performed the experiments with academic and technical assistance from OSY, KK, TY, TF, SU, and HM. KS, KK, and HM analyzed the data and wrote the first draft of the manuscript. All authors critically reviewed the manuscript and approved the final submission.

## Acknowledgements

The authors would like to express our gratitude to emeritus professor Hiroshi Chiura of Tokyo University of Agriculture and Technology for giving us helpful advice.

## Supplementary material

