## Supplemental Data 1-4, and Certificate for "Novel endornaviruses infecting *Phytophthora cactorum* that attenuate vegetative growth, promote sporangia formation, and confer hypervirulence to the host oomycete": Certicaate for proofing. Sakuta Moriyama.pdf

NAI
2-21-1 Tsuruya-cho, Kanagawa-Ku, Yokohama, Kanagawa Pref. 221-0835 Japan

2024/11/15

To Whom It May Concern:

An experienced editor whose first language is English has carefully reviewed this abstract entitled:

**Novel endornaviruses infecting *Phytophthora cactorum* that attenuate vegetative growth, promote sporangia formation, and confer hypervirulence to the host oomycete.**

Our company specializes in scientific editing for papers written by scientists whose native language is not English.

Shuji Ito
President NAI, Inc.
