## Supplemental Data 1-4, and Certificate for "Novel endornaviruses infecting *Phytophthora cactorum* that attenuate vegetative growth, promote sporangia formation, and confer hypervirulence to the host oomycete": Supplemental Data.pdf

**Amino acid(similarity)**

**Nucleotide**

| | PcAEV1 | PcAEV2 | PcAEV3 | PcAEV4 | PcAEV5 |
|---|---|---|---|---|---|
| PcAEV1 | | 20.1%(35.0%) | 19.3%(33.8%) | 19.9%(34.0%) | 19.2%(32.5%) |
| PcAEV2 | 44.8% | | 44.6%(63.0 %) | 58.4%(75.4 %) | 44.4%(63.3 %) |
| PcAEV3 | 44.6% | 54.0% | | 44.8%(63.1 %) | 83.7%(91.9 %) |
| PcAEV4 | 44.6% | 59.6% | 53.4% | | 45.7%(63.7 %) |
| PcAEV5 | 44.3% | 54.0% | 73.2% | 54.6% | |

| | | | | |
|---|---|---|---|---|
| ~ 20 % | 21 ~ 40 % | 41 ~ 60 % | 61 ~ 80 % | 81 % ~ |

**Supplemental Figure S1. Pairwise sequence alignment (PSA) comparing the complete nucleotide and amino acid sequences of PcAEV1-5.** PSA was performed using EMBOSS Needle, and the identities (similarities in parentheses) of the complete nucleotide and amino acid sequences are shown. To clarify the results, darker blue indicates higher percentages, with 100% identity removed by the diagonal line.

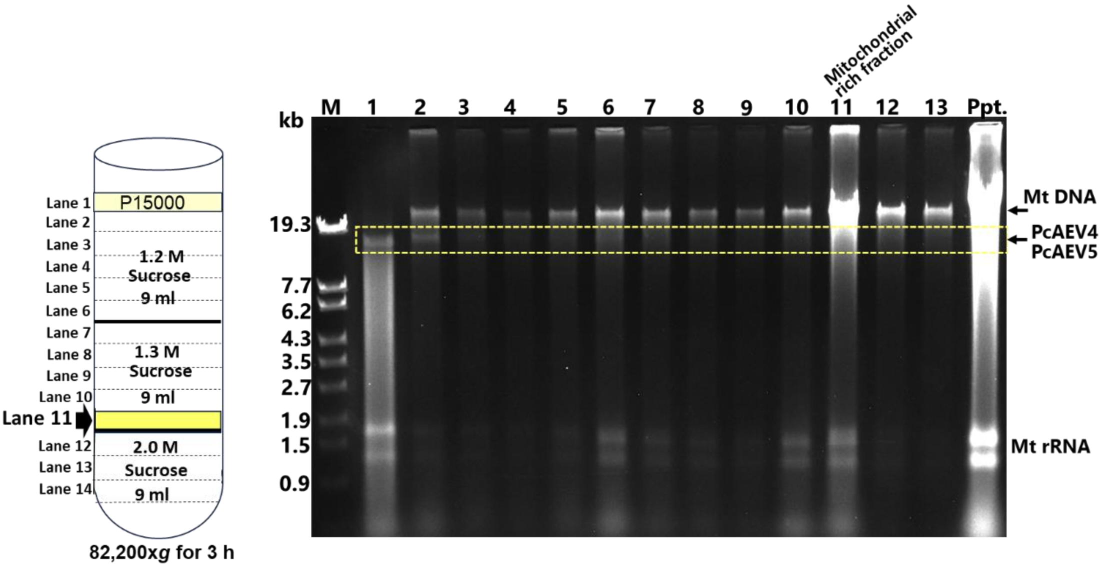


**Supplemental Figure S2. Discontinuous sucrose gradient fractionation of crude mitochondrial fraction P15000.** Fractions obtained were labeled 1 through 14 from top to bottom (Lanes 1-14). Total nucleic acids were extracted from 1/10 of each fraction and subjected to agarose gel electrophoresis. Lane M: DNA marker (250 ng of λDNA digested with EcoT14I). The positions of PcAEV4 and PcAEV5 dsRNA are indicated by yellow dashed lines and arrows. Mt DNA: mitochondrial genomic DNA; Mt rRNA: mitochondrial ribosomal RNA.

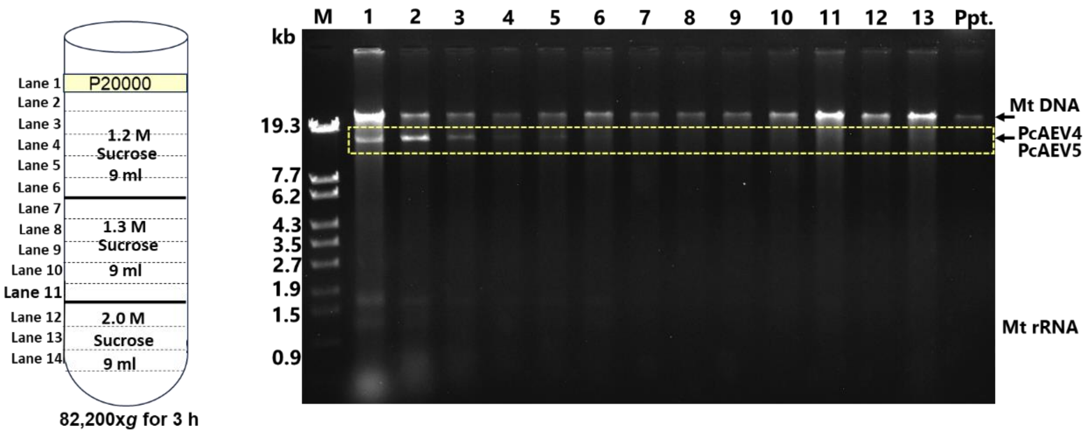


**Supplemental Figure S3. Discontinuous sucrose gradient fractionation of P20000.** Fractions obtained were labeled 1 through 14 from top to bottom (Lanes 1-14). Total nucleic acids were extracted from 1/10 of each fraction and subjected to agarose gel electrophoresis. Lane M: DNA marker (250 ng of λDNA digested with EcoT14I). The positions of PcAEV4 and PcAEV5 dsRNA are indicated by yellow dashed lines and arrows. Mt DNA: mitochondrial genomic DNA; Mt rRNA: mitochondrial ribosomal RNA.

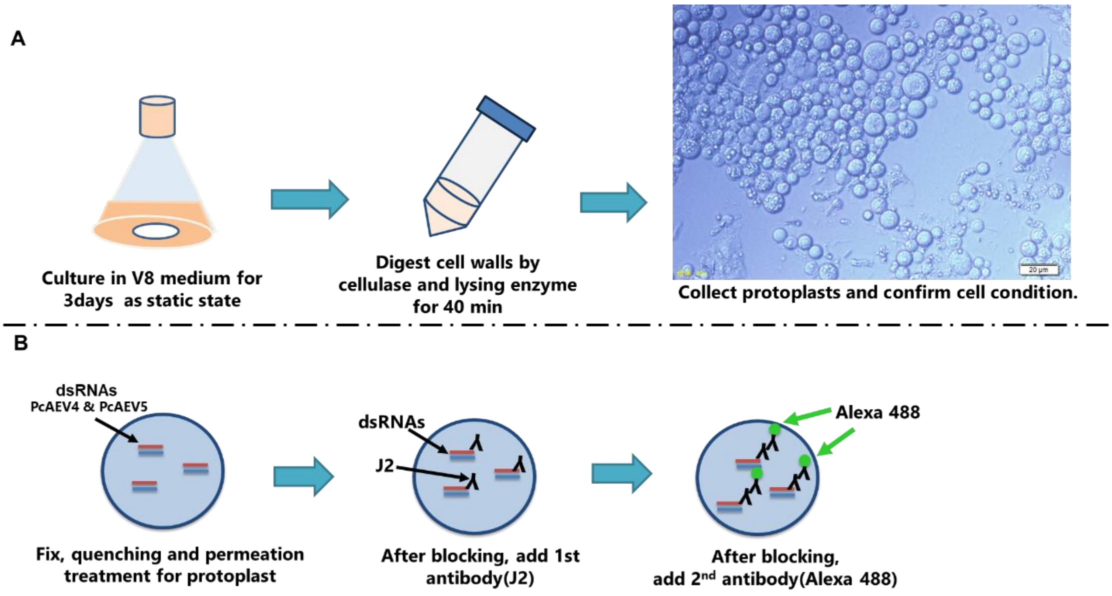


**Supplemental Figure S4. IFA scheme for detecting dsRNA in *Phytophthora* spp. cells. (A)** Schematic of protoplast preparation for IFA. **(B)** Schematic of IFA in protoplasts.
